# Aneuploidy sensitizes cells to SREBP-pathway inhibition in squamous cell carcinoma

**DOI:** 10.64898/2026.05.04.722276

**Authors:** Nadja Zhakula-Kostadinova, Sejal Jain, Zeinab Amini-Farsani, Jiankang Zhang, Mari Nakamura, Laura Byron, Joan J. Castellano Perez, Chloe Paolucci, Rohan Munoth, Fereshteh Zandkarimi, Yuka Takemon, Marco Marra, Brian Henick, Anjali Saqi, Tannishtha Reya, Matthew Meyerson, Alison M. Taylor

**Affiliations:** Department of Genetics and Development, Columbia University Vagelos College of Physicians and Surgeons; Department of Medical Oncology, Dana-Farber Cancer Institute; Department of Pathology and Cell Biology, Herbert Irving Comprehensive Cancer Center, Columbia University Vagelos College of Physicians and Surgeons; Department of Chemistry, Columbia University; Michael Smith Laboratories, University of British Columbia, Vancouver, Canada; Department of Medical Genetics, Faculty of Medicine, University of British Columbia, Vancouver, Canada; Department of Medical Oncology, Columbia University Medical Center; Department of Genetics and Medicine, Harvard Medical School; Broad Institute of Harvard and MIT

**Author notes:** **Corresponding Author:** Alison M. Taylor, Columbia University Irving Cancer Research Center, 1130 St. Nicholas Ave, Rm 301B, New York, NY 10032.

## Abstract

Squamous cell carcinomas (SCCs) in the lung, head and neck, cervix, and esophagus are characterized by widespread chromosome-arm aneuploidies, most frequently recurrent 3q-gain. However, how these alterations influence cancer development and therapeutic vulnerabilities remains unclear. To identify aneuploidy-driven therapeutic targets, we performed genome-wide CRISPR interference (CRISPRi) and drug-repurposing screens in isogenic immortalized lung epithelial cells harboring chromosome 3-disomy or 3q-gain. Both screens converged on a mevalonate pathway dependency specific to 3q-gain cells, which exhibited heightened sensitivity to sterol regulatory element-binding protein (SREBP) disruption. Rescue experiments demonstrated that these vulnerabilities were on target and that pathway inhibition preferentially causes apoptosis in 3q-gain cells. Transcriptomic and lipidomic profiling revealed 3q-gain-associated alterations in SREBP activation, cholesterol and fatty-acid biosynthesis, and lipid composition. Perturbing SREBP signaling impaired viability in SCC cell lines and suppressed tumor growth in xenografts with 3q-gain. These findings identify an aneuploidy-driven, targetable vulnerability in SCC.

**Significance:** Here, we demonstrate that SCC-recurrent 3q-gain is a selective vulnerability to SREBP-pathway inhibition. We identify an aneuploidy-driven therapeutic liability in squamous tumors for lipid-targeted precision therapies, providing a framework for targeted treatment in SCC.

## Introduction

Squamous cell carcinomas (SCCs) arise from stratified epithelial tissues, encompassing malignancies of the lung, head and neck, esophagus, cervix, and other anatomic sites (1,2). Human papillomavirus (HPV) is associated with approximately 25-30% of head and neck SCCs and over 90% of cervical SCCs (3,4). HPV-negative SCCs are associated with particularly poor survival outcomes and mortality rates approaching 80% across disease sites (5). Despite their aggressive clinical behavior, SCCs remain largely devoid of canonical, targetable oncogenic drivers that define other epithelial cancers, such as *EGFR*, *ALK*, or *KRAS* alterations in adenocarcinoma (5,6). Instead, SCCs frequent inactivation of tumor suppressors such as *TP53*, *PTEN*, and *CDKN2A*, alongside recurrent amplification of lineage-defining transcription factors including *TP63* and *SOX2* (7,8). Notably, SCC genomes, regardless of HPV status, are characterized by extensive copy-number alterations and chromosome arm aneuploidy–defined as gains or losses of whole chromosomes or chromosome arms (9).

Aneuploidy is a hallmark of cancer and a pervasive feature of solid tumors (10). Although aneuploidy is generally poorly tolerated in normal cells (11,12), chromosome arm aneuploidies are the most frequent somatic alteration in cancer, with approximately 90% of solid tumors harboring at least one arm-level copy-number alteration (13). Importantly, chromosome arm aneuploidies recur in tumor-type-specific patterns, clustering according to tissue lineage and histology (13). In SCCs, chromosome 3 alterations—specifically 3p-deletion and 3q-gain—are the most lineage-defining and recurrent copy-number events (4,13,14). These alterations are detectable in premalignant lesions and are considered early drivers of squamous tumorigenesis (15–18). Chromosome 3q-gain distinguishes lung SCC from lung adenocarcinoma and results in overexpression of key squamous lineage regulators *SOX2* and *TP63* (18). However, the broader consequences of gain of the whole 3q arm remain unclear. Recurrent chromosome arm alterations likely exert effects that extend beyond cancer-promoting drivers and could serve as therapeutic targets (19–22). Thus, identifying the functional dependencies created by lineage-defining aneuploidies may uncover therapeutically actionable vulnerabilities in SCC.

Here, we define dependencies conferred by 3q-gain using an isogenic human lung epithelial model that recapitulates the putative cell-of-origin of lung SCC (13). Through genome-wide CRISPR interference (CRISPRi) and drug repurposing screens, we identify a selective sensitivity of 3q-gain cells to inhibition of the mevalonate pathway, revealing a dependency on sterol regulatory element–binding protein 1 (SREBP1) and its downstream metabolic target, the rate-limiting enzyme HMG-CoA reductase (HMGCR). We demonstrate that 3q-gain cells display a lipid and transcriptional state characterized by attenuated SREBP activation in response to sterol depletion. We further identify chromosome 3q-encoded regulators of ER-to-Golgi trafficking as modulators of mevalonate pathway sensitivity. Finally, we show that these vulnerabilities are conserved in cancer through *in silico* genetic interaction analyses of CRISPR dependency datasets and *in vivo* SCC xenograft models. Collectively, these findings illustrate how a recurrent chromosome arm aneuploidy can expose therapeutically actionable dependencies in squamous malignancies.

## Results

### Chromosome 3q-gain correlates with genetic dependency on SREBF1

Given the high prevalence of 3q-gain in squamous malignancies and its early emergence in disease evolution, we sought to define vulnerabilities created by this aneuploidy in a non-transformed epithelial context. We used isolated isogenic cell lines that harbor 3q-disomy (wild-type; WT) or 3q trisomy (3q-gain) (13,23). These cells were engineered in lung basal epithelial cells, the putative cell of origin for lung SCC (24). The lung basal epithelial cells were previously immortalized with Simian Virus 40 (SV40) T antigen and telomerase reverse transcriptase (TERT) (24), functionally inactivating the p53 and RB pathways. We transduced WT and 3q-gain cells with a pooled, barcoded genome-wide CRISPRi library to identify genotype-specific genetic dependencies (**Figure 1A**) (25). After fourteen days of culture with selection, we quantified sgRNA representation using MAGeCK. The most significantly differentially depleted sgRNAs in 3q-gain cells targeted sterol regulatory element–binding transcription factor 1 (*SREBF1)* (log_2_FC = -1.43, *p* = 2.6 × 10⁻, **Figure 1B**) which encodes sterol regulatory element-binding protein 1 (SREBP1), a transcriptional regulator of lipid metabolic programs (26). Individual *SREBF1* knockdown with three CRISPRi guides (**Supplementary Figure S1A**) significantly reduced viability preferentially in 3q-gain cells in 2/3 guides tested (*p* = 0.05 and 0.04, **Figure 1C**), confirming the screen result.

**Figure 1.**
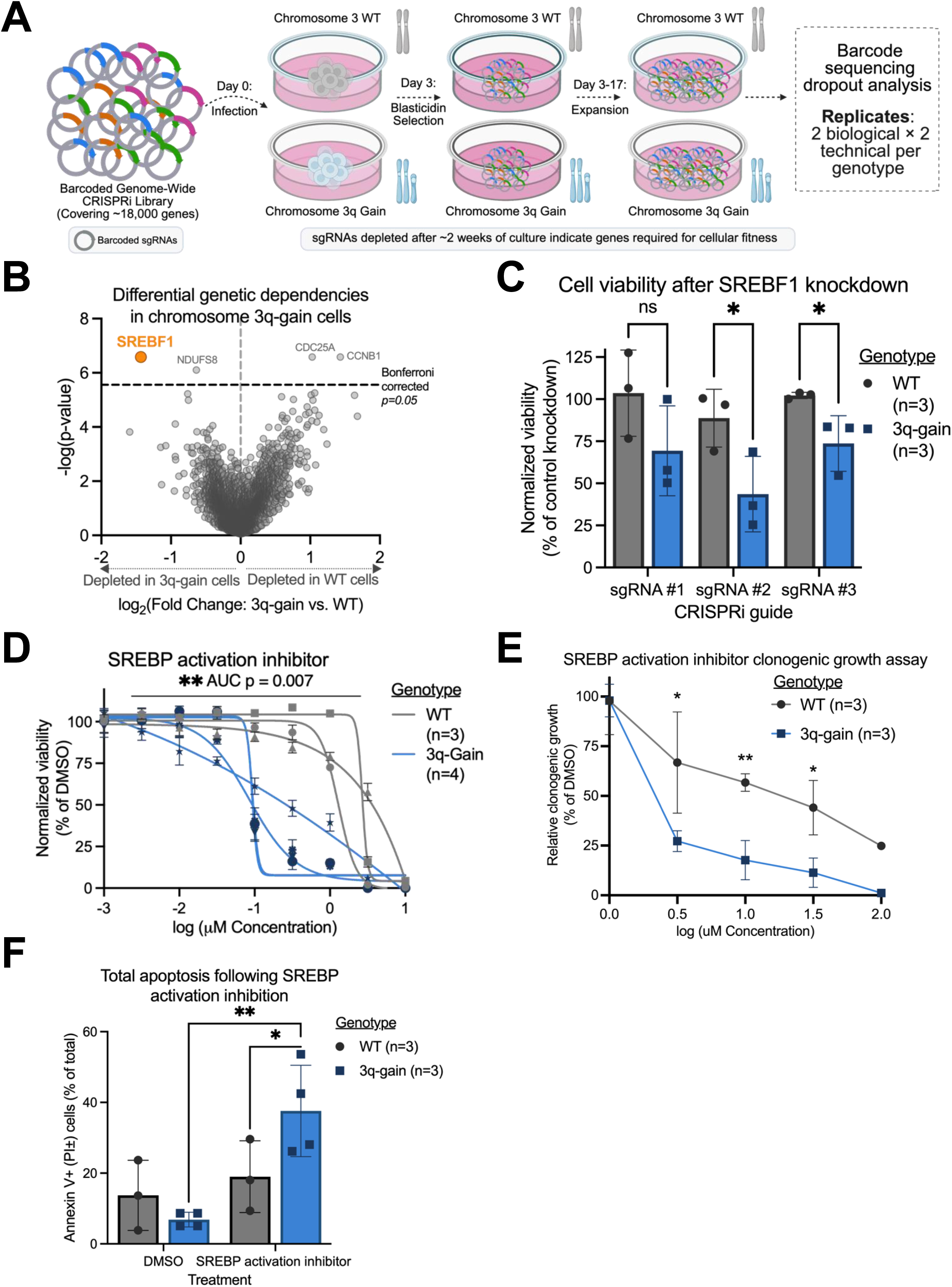
Genome-wide CRISPRi screening identifies genotype-specific genetic dependencies in 3q-gain cells. **(A)** Schematic of pooled, barcoded genome-wide CRISPR interference (CRISPRi) screening strategy performed in isogenic WT and 3q-gain cells. A barcoded genome-wide CRISPRi library targeting ∼18,000 genes was introduced into chromosome 3 wild-type (WT) and 3q-gain human airway lung epithelial cells. Following lentiviral transduction, cells were subjected to blasticidin selection beginning 72 hours post-infection and expanded for 2 weeks prior to endpoint collection. sgRNA abundance was quantified by sequencing, with 2 biological and 2 technical replicates used per genotype. Created in BioRender. **(B)** Differential genetic dependencies between WT and 3q-gain cells identified by CRISPRi screening. Each point represents sgRNAs targeting a single gene, plotted by log₂ fold change in dependency (3q-gain vs WT) and −log₁₀(p-value). sgRNAs significantly depleted and enriched in 3q-gain cells relative to WT are highlighted in orange (false discovery rate (FDR)-adjusted p < 0.05), with negative fold changes indicating stronger depletion in 3q-gain cells. **(C)** Cell viability following CRISPRi-mediated knockdown of SREBF1 using three independent sgRNAs in WT and 3q-gain cells. Viability was measured by CellTiter-Glo 5 days after selection and normalized to non-targeting control sgRNAs. Each dot represents an independent clone (n = 3 WT, n = 3 3q-gain). Statistical significance was assessed by two-way ANOVA with Tukey’s multiple comparison test. **(D)** Dose-response of WT and 3q-gain cells treated with the SREBP activation inhibitor fatostatin. Cell viability was measured by CellTiter-Glo 72 h after treatment and normalized to DMSO-treated controls. Each line represents an independent clone (WT, *n* = 3; 3q-gain, *n* = 4), with each data point reflecting the mean of three technical replicates. Area under the curve (AUC) values were calculated for each clone and compared between genotypes using a two-sided Student’s t-test. **(E)** Quantification of clonogenic growth from **Supplementary Figure 1B**, normalized to DMSO-treated controls. Each point represents the mean relative clonogenic growth across independent clones (WT, *n* = 3; 3q-gain, *n* = 3). Statistical significance assessed by Student’s t-test comparing genotypes at each concentration. **(F)** Total apoptotic cell death following 72 h of treatment with the SREBP activation inhibitor fatostatin (10μM), measured by Annexin V and propidium iodide (PI) staining. Bars represent percentage of Annexin V⁺/PI⁺ and Annexin V⁺/PI- cells. Statistical significance assessed by two-tailed unpaired Student’s *t* tests comparing genotypes at each concentration. Bars indicate mean ± s.d. P values are represented as: *, P < 0.05; **, P < 0.01; ***, P < 0.001; ****, P < 0.0001.

We next examined whether chemical inhibition of SREBP activation phenocopied the genetic dependency observed in the screen. Treatment with fatostatin, an inhibitor of SREBP activation, preferentially reduced viability in 3q-gain cells in a dose-dependent manner (AUC FC = 0.649, *p* = 0.007, **Figure 1D**). This differential sensitivity was also evident in a two-week growth assay, where inhibition of SREBP activation was significantly more toxic to 3q-gain cells than WT counterparts (**Figure 1E, Supplementary Figure S1B**). Consistent with these growth defects, SREBP inhibition induced significantly higher levels of total apoptotic cell death in 3q-gain cells, measured by Annexin V and propidium iodide staining (*p* = 0.029, **Figure 1F, Supplementary Figure S1C-D**). Together, these data demonstrate that chromosome 3q-gain correlates with genetic and chemical vulnerabilities to SREBP inhibition in lung epithelial cells.

### Chromosome 3q-gain correlates with increased sensitivity to HMGCR inhibition

To determine whether chromosome 3q-gain also correlated with differential sensitivity to chemical perturbations, we screened 5439 compounds from a drug-repurposing library (27), comparing viability between WT and 3q-gain cells after 72 hours of treatment at 2.5µM (**Supplementary Figure S2A**). After two rounds of validation in multiple clones per genotype, the most prominent genotype-selective hit was mevastatin (**Figure 2A**). Mevastatin, the first discovered statin, belongs to a class of drugs that inhibit 3-hydroxy-3-methylglutaryl-CoA reductase (HMGCR), the rate-limiting enzyme in the mevalonate pathway (**Figure 2B**) (28,29).

**Figure 2.**
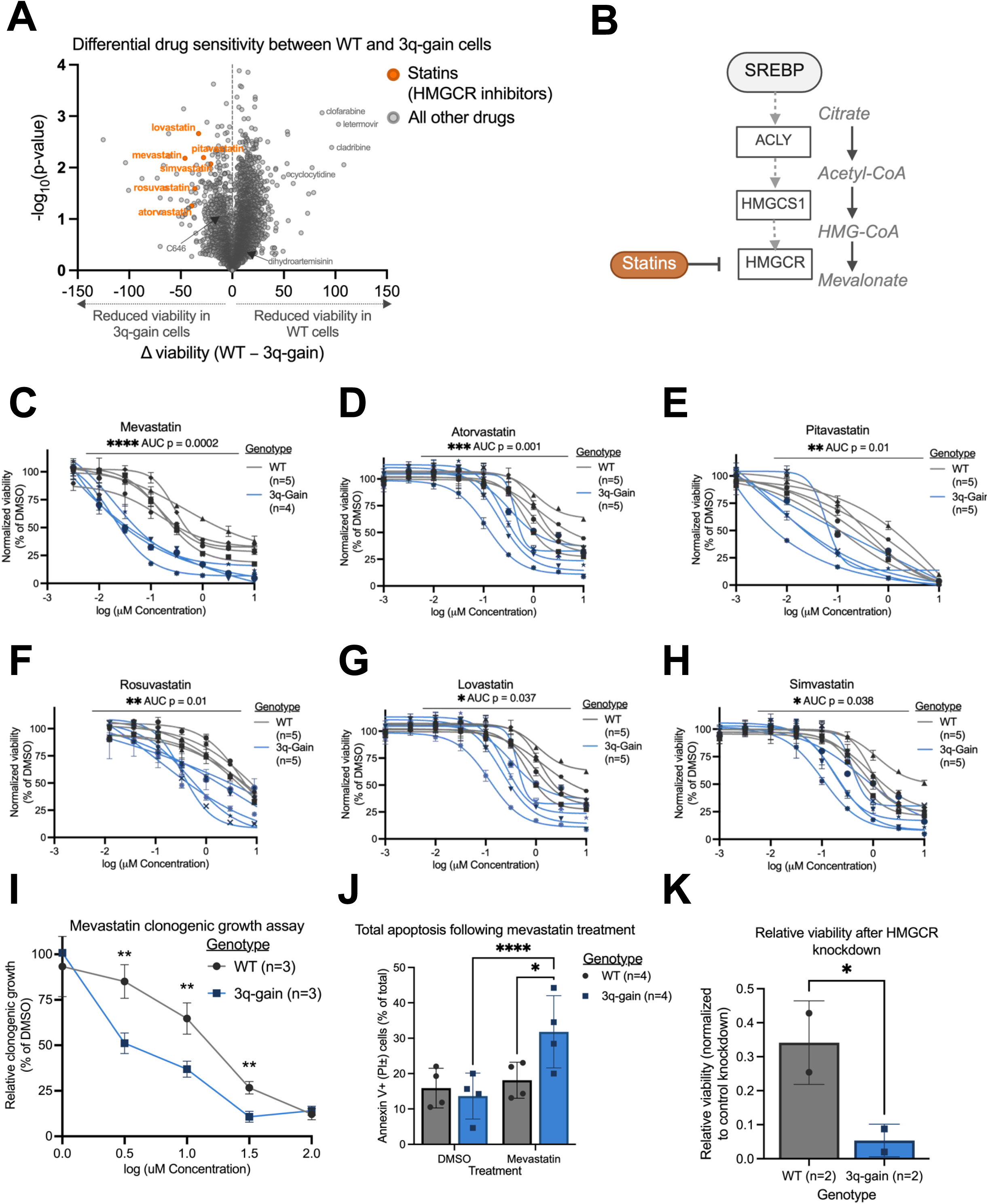
Pharmacologic screening identifies selective sensitivity of 3q-gain cells to HMGCR inhibition. **(A)** Differential drug sensitivity between WT and 3q-gain cells following treatment with a small-molecule drug-repurposing library at 2.5μM. Each point represents a compound, plotted by the difference in normalized viability between genotypes (WT−3q-gain) and −log₁₀(p-value). Named compounds were validated by individual dose curves, with statins in orange. **(B)** Schematic overview of the mevalonate pathway, highlighting the position of 3-hydroxy-3-methylglutaryl–CoA reductase (HMGCR) and its regulation by SREBPs. Statins inhibit HMGCR, a rate-limiting enzyme of cholesterol biosynthesis. Created in BioRender. **(C-H)** Dose–response analysis of WT and 3q-gain AALE cells treated with individual statins, including mevastatin (**C**), simvastatin (**D**), pitavastatin (**E**), lovastatin (**F**), atorvastatin (**G**), and rosuvastatin (**H**). CellTiter-Glo luminescence 72 h after treatment and normalized to DMSO-treated controls. Each line represents an independent clone, with each data point reflecting the mean of three technical replicates. AUC values were calculated for each clone and compared between genotypes using a two-sided Student’s t-test. **(I)** Quantification of clonogenic growth from **Supplementary Figure S2B**, normalized to DMSO-treated controls. Each point represents the mean relative clonogenic growth of an independent clone. Statistical significance at individual doses was assessed by two-sided Student’s t-test comparing genotypes at each concentration. **(J)** Total apoptotic cell death following 72 h of mevastatin treatment (10μM), measured by Annexin V and propidium iodide (PI) staining. Bars represent the percentage of Annexin V⁺ and/or PI⁺ cells. Each dot represents an independent clone. Statistical significance was assessed by two-way ANOVA with Tukey’s multiple comparison test. **(K)** CRISPRi-mediated knockdown of HMGCR at day 20 in WT and 3q-gain cells. Cell viability is normalized to control knockdown within each genotype. Bars indicate mean ± s.d. *P* value thresholds are defined in Figure 1.

Statins consistently ranked among the compounds with the largest differential effects. Of the 21 statins present in the drug-repurposing library, 8 exhibited differential activity in the primary screen. Follow-up validation with full dose curves confirmed increased sensitivity of 3q-gain cells to 6 statins spanning distinct chemical classes, including both lipophilic and hydrophilic agents: mevastatin (area under the curve fold change [AUC FC] = 0.576, *p* = 0.0002, **Figure 2C**), atorvastatin (AUC FC = 0.614, *p* = 0.001 (**Figure 2D**), pitavastatin (AUC FC = 0.694, *p* = 0.01; **Figure 2E**), rosuvastatin (AUC FC = 0.789, *p* = 0.01; **Figure 2F**), lovastatin (AUC FC = 0.841, *p* = 0.037; **Figure 2G**), and simvastatin (AUC FC = 0.855, *p* = 0.038; **Figure 2H**). Notably, mevastatin exhibited the strongest differential effect between WT and 3q-gain cells (**Figure 2C**) and was therefore selected for subsequent experiments.

Consistent with short-term viability measurements, treatment with mevastatin significantly impaired long-term clonogenic growth of 3q-gain cells compared to WT counterparts (**Figure 2I, Supplementary Figure S2B**) and had a significant reduction in absolute cell number (**Supplementary Figure S2C**). In parallel, mevastatin treatment also induced significantly higher levels of total apoptotic cell death in 3q-gain cells, measured by Annexin V and propidium iodide staining (*p* = 0.01, **Figure 2J, Supplementary Figure S2D**) and caspase-3/7 activation (*p* < 0.0001, **Supplementary Figure S2E**). CRISPRi-mediated knockdown of *HMGCR* also caused a significant reduction of viability in 3q-gain cells relative to WT cells (*p* = 0.01, **Figure 2K, Supplementary Figure S2F**). Together, these data demonstrate that our cells with 3q-gain are more vulnerable to chemical and genetic inhibition of HMGCR.

### Dependence of 3q-gain cells on mevalonate pathway flux and SREBP regulation

SREBPs coordinate transcriptional output of the mevalonate pathway (schematic in **Figure 3A**) (26,30). To determine whether pathway sensitivity in 3q-gain cells reflects on-target disruption of the mevalonate pathway, we first asked whether mevalonate could rescue statin-induced cell death. Supplementation with exogenous mevalonate significantly attenuated mevastatin-induced apoptosis in 3q-gain cells, restoring viability to levels comparable to WT controls (*p* = 0.001, **Figure 3B, Supplementary Figure S3A**). Consistent with this effect on cell death, mevalonate also rescued viability following mevastatin treatment (*p* = 0.008, **Figure 3C**), demonstrating that statin sensitivity in 3q-gain cells is driven by perturbation of the mevalonate pathway.

**Figure 3.**
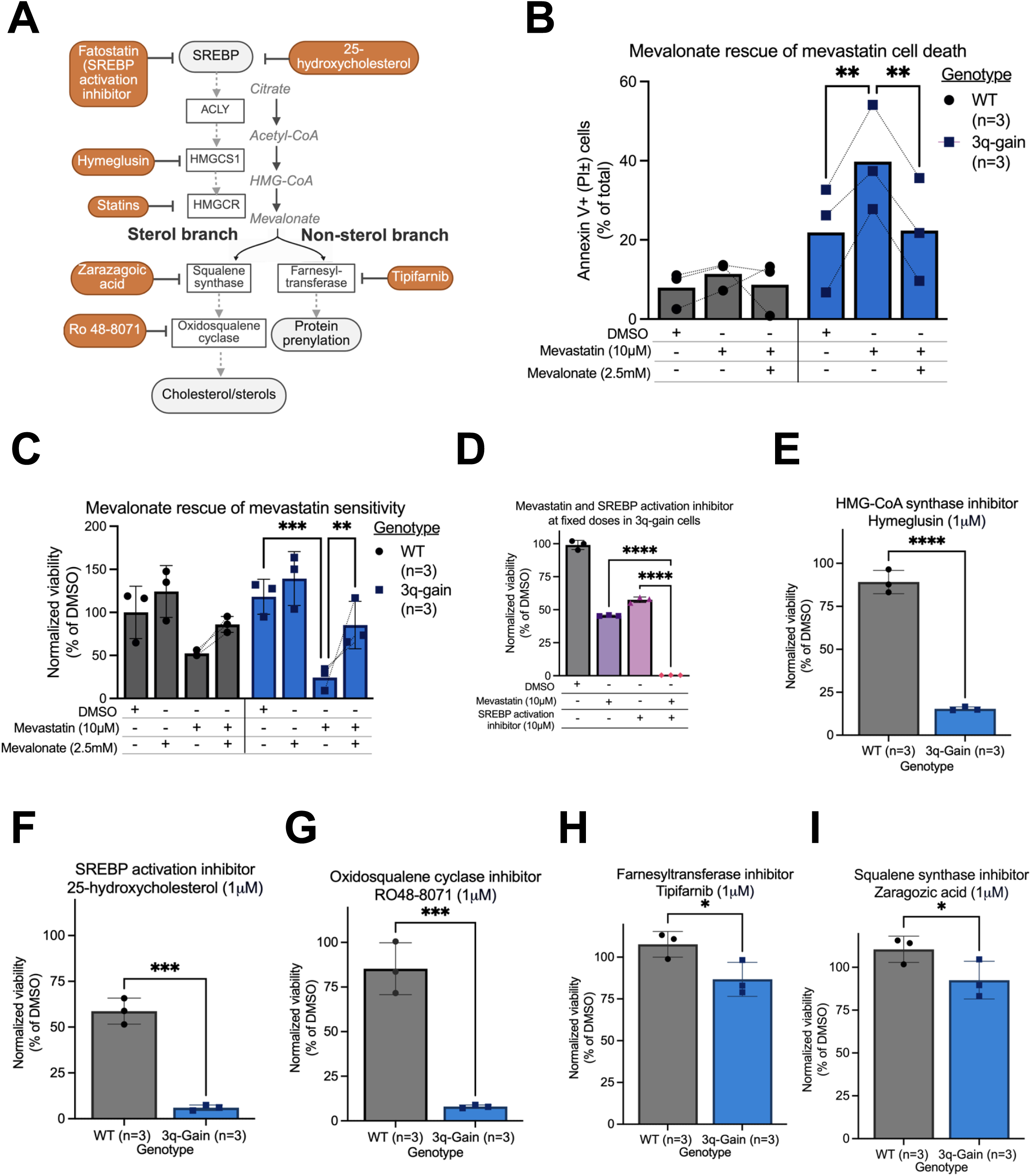
Sensitivity of 3q-gain cells reflects dependence on mevalonate pathway. **(A)** Schematic overview of the mevalonate pathway, highlighting sterol and non-sterol branches and sites of genetic or pharmacologic perturbation used in this study. Created in BioRender. **(B)** Annexin V/PI staining in WT and 3q-gain cells following 72 h of mevastatin and/or mevalonate treatment. Total apoptosis is represented by Annexin V^+^/PI^-^ and Annexin V^+^/PI^+^ populations. Individual clones across treatments are represented by dotted lines. **(C)** Cell viability following 72 h of mevastatin and/or mevalonate treatment in WT and 3q-gain cells, as measured by CellTiter-Glo. **(D)** Cell viability following combined treatment with mevastatin (HMGCR inhibition) and the SREBP activation inhibitor fatostatin at a fixed, representative dose in WT and 3q-gain cells. **(E-I)** Pharmacologic perturbation of non-statin nodes within the mevalonate pathway. Shown are inhibitors of HMG-CoA synthase (hymeglusin; **E**), SREBP activation (25-hydroxycholesterol; **F**), oxidosqualene cyclase (RO48-8071; **G**), farnesyltransferase (tipifarnib; **H**), and squalene synthase (zaragozic acid; **I**). Data are shown as mean ± SEM. Unless otherwise indicated, each dot represents a biologically independent clone, with data points reflecting the mean of three technical replicates. Statistical analyses were performed using two-way ANOVA with Sidak’s multiple-comparisons test for genotype–treatment comparisons (**B-C**) and two-sided paired t-tests for pairwise comparisons between genotypes (**D-I**). *P* value thresholds are defined in Figure 1.

Since HMGCR activity and SREBP-dependent transcription converge on the mevalonate pathway, we next asked whether combined inhibition would further exacerbate growth defects in 3q-gain cells. Co-treatment with mevastatin and the SREBP activation inhibitor fatostatin resulted in a significantly greater reduction in viability than either agent alone in 3q-gain cells (*p* < 0.0001, **Figure 3D**), consistent with a synergistic interaction between inhibition of HMGCR and disruption of SREBP-dependent transcription. We quantified synergy with Highest Single Agent (HSA) analysis, which assesses whether the observed combination effect exceeds that of the most effective single agent at matched doses (HSA score = 12.11, *p* = 5.98 × 10⁻¹², **Supplementary Figure S3B)**.

To further delineate which nodes within the mevalonate pathway contribute to the selective vulnerability of 3q-gain cells, we tested inhibitors targeting distinct enzymatic steps and regulatory mechanisms within the pathway. Inhibition of HMG-CoA synthase 1 (HMGCS1) using hymeglusin, which blocks conversion of acetyl-CoA to HMG-CoA upstream of HMGCR, preferentially reduced viability in 3q-gain cells compared to WT cells (*p* < 0.0001 **Figure 3E**). Similarly, CRISPRi-mediated knockdown of HMGCS1 significantly reduced viability in 3q-gain cells relative to WT (*p* = 0.0087, **Supplementary Figure S3C-D**). Inhibition of SREBP activation using 25-hydroxycholesterol, which suppresses SREBP processing through sterol-mediated feedback, significantly impaired viability in 3q-gain cells (*p* = 0.0001, **Figure 3F**). Inhibition of the sterol branch of the mevalonate pathway with RO48-8071, an oxidosqualene cyclase inhibitor, similarly differentially reduced viability in 3q-gain cells (*p* = 0.0004, **Figure 3G**). Inhibitors targeting downstream branches of the mevalonate pathway, including farnesyltransferase inhibition with tipifarnib and squalene synthase inhibition with zaragozic acid, likewise reduced viability more strongly in 3q-gain cells than in WT cells, but with quantitatively weaker genotype-specific effects relative to upstream enzymatic inhibition and SREBP regulation (*p* = 0.022 and 0.04, respectively, **Figure 3H-I**). Together, these data demonstrate that the heightened vulnerability in our 3q-gain cells reflects a dependence on the mevalonate pathway and its upstream transcriptional regulation by SREBP.

### Chromosome 3q-gain is associated with a distinct lipid and transcriptional state

As 3q-gain cells are more sensitive to mevalonate pathway inhibition, we next characterized lipid homeostasis in isogenic WT and 3q-gain cells. 3q-gain cells exhibited reduced lipid storage (measured by BODIPY 493/503 staining) relative to WT cells (*p*= 0.044, **Figure 4A**) which was further accentuated with mevastatin treatment (*p* = 0.0003, **Figure 4A**). Consistent with altered lipid storage, direct measurement of cholesterol pools revealed genotype-specific differences in cholesterol handling across conditions. Intracellular free and total cholesterol levels were reduced in 3q-gain cells relative to WT cells under basal conditions (*p* = 0.04 and 0.003 respectively) and following mevastatin treatment (*p* = 0.0025 and 0.001, respectively, **Figure 4B, Supplementary Figure S4A**). Intracellular levels were restored by supplementation with exogenous mevalonate (*p* < 0.0001, **Figure 4B**, *p* = 0.001, **Supplementary Figure S4A**). In contrast, extracellular free and total cholesterol levels were significantly elevated in 3q-gain cells compared to WT cells (*p* = 0.03 and 0.02), again enhanced with mevastatin treatment (*p* < 0.0001, **Figure 4C**, *p* = 0.0004, **Supplementary Figure S4B**). To assess whether these differences reflected altered cholesterol acquisition, we measured basal cholesterol uptake capacity. 3q-gain cells exhibited significantly reduced cholesterol uptake relative to WT cells (*p* = 0.018, **Figure 4D**), consistent with impaired maintenance of intracellular cholesterol pools.

**Figure 4.**
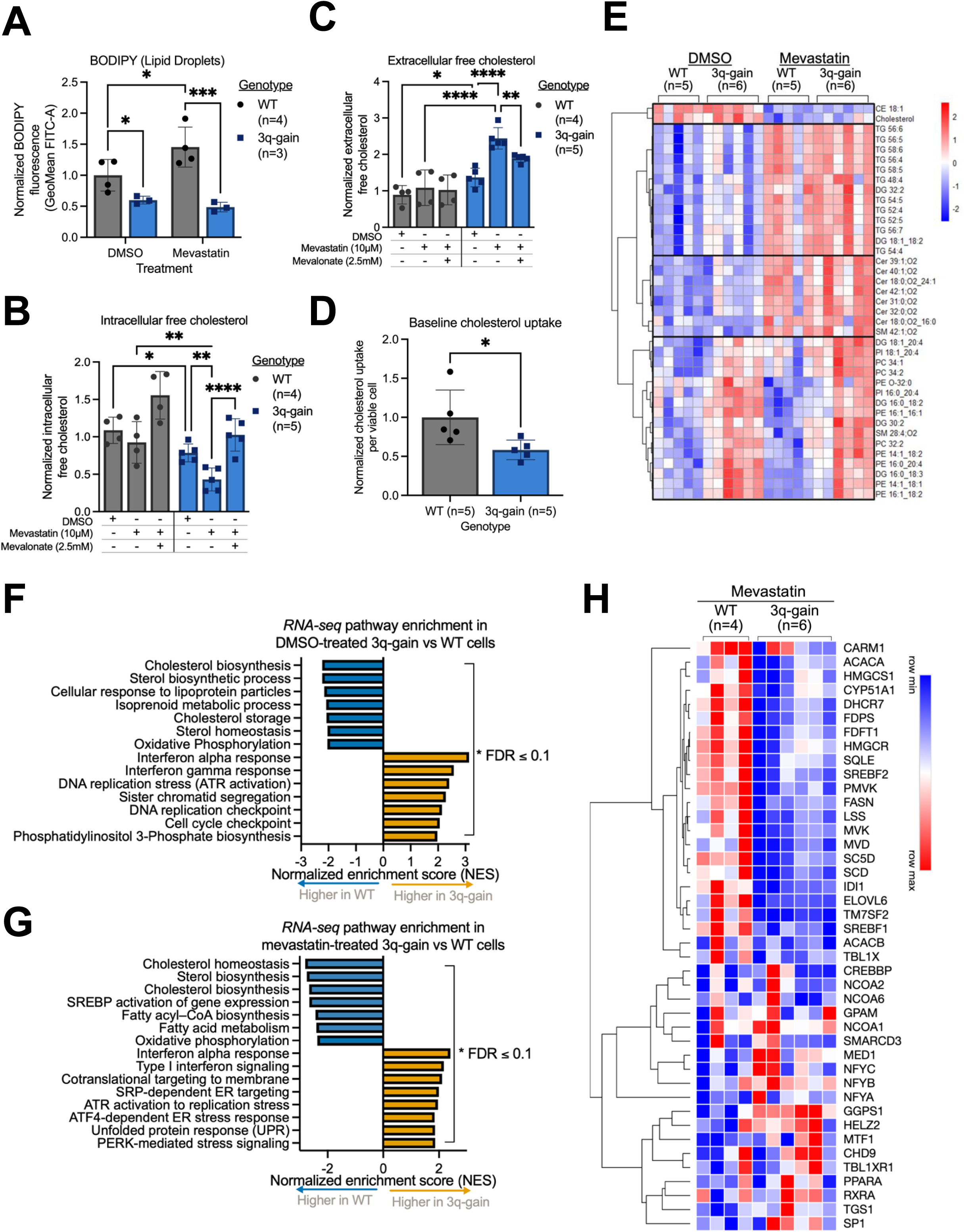
Chromosome 3q-gain is associated with a distinct lipid and transcriptional state. **(A)** Neutral lipid accumulation assessed by BODIPY staining and quantified as fluorescence intensity by flow cytometry in WT and 3q-gain cells following treatment with DMSO or mevastatin (10 µM). **(B)** Intracellular free cholesterol levels in WT and 3q-gain cells treated with mevastatin (10 µM) with or without mevalonate supplementation (2.5 mM), measured by the Cholesterol-Glo assay and normalized to WT DMSO-treated controls. **(C)** Extracellular free cholesterol levels under the same treatment conditions as in (**B**). **(D)** Baseline cholesterol uptake in WT and 3q-gain cells under steady-state conditions, measured using a fluorescent NBD-cholesterol uptake assay. **(E)** Heatmap showing relative abundance of cholesterol and lipid species in WT and 3q-gain cells treated with DMSO or mevastatin, as measured by untargeted lipidomics. Lipid species are grouped by class, including triglycerides (TG), diglycerides (DG), phosphatidylcholines (PC), phosphatidylethanolamines (PE), phosphatidylinositols (PI), ceramides (Cer), sphingomyelins (SM), and cholesterol esters (CE). **(F)** Gene set enrichment analysis (GSEA) of RNA-seq data showing pathways enriched in DMSO-treated 3q-gain versus WT cells. **(G)** GSEA of RNA-seq data showing pathways enriched in mevastatin-treated 3q-gain versus WT cells following 72 h of treatment. **(H)** Heatmap of selected cholesterol biosynthesis and SREBP1-regulated transcriptional targets derived from the RNA-seq dataset shown in **G**. Data are shown as mean ± SEM. Dots represent independent clones as biological replicates. Statistical significance was assessed using two-way ANOVA with Tukey’s multiple comparison test for panels **A–C**, two-tailed unpaired Student’s *t* tests for panel **D**, and one-way ANOVA with FDR < 0.05 for panel **E**. GSEA and pathway enrichment analyses were performed using pre-ranked gene lists with an FDR ≤ 0.1. Normalized enrichment scores (NES) are shown. *P* value thresholds are defined in Figure 1.

We next performed lipidomic profiling with and without mevalonate pathway inhibition to capture broader alterations in lipid composition associated with 3q-gain (**Supplementary Table S1**). Mevastatin treatment was associated with a significant reduction in free cholesterol and cholesterol esters across genotypes, consistent with on-target inhibition of cholesterol synthesis (**Figure 4E**). Mevastatin-treated cells exhibited increased abundance of triglycerides and ceramides, reflecting coordinated changes in lipid species following mevalonate pathway perturbation. In contrast, chromosome 3q-gain cells displayed a distinct lipidomic signature independent of treatment with increased levels of multiple phospholipid species, including phosphatidylethanolamines (PEs), phosphatidylinositols (PIs), and phosphatidylcholines (PCs), as well as elevated diacylglycerides (DGs), relative to WT cells (**Figure 4E**). Together, these data suggest that chromosome 3q-gain is associated with distinct patterns of lipid composition.

To assess transcriptional phenotypes, we performed RNA-seq of WT and 3q-gain cells with vehicle, mevastatin, or fatostatin treatment (**Supplementary Table S2**). Pathways related to cholesterol biosynthesis and SREBP-regulated gene expression were significantly lower in 3q-gain cells relative to WT cells under basal conditions, alongside increased expression of interferon and DNA replication pathways (FDR ≤ 0.1, **Figure 4F**). Following mevastatin treatment, 3q-gain cells exhibited lower expression of SREBP1 target genes and cholesterol and fatty acid synthesis pathways compared to WT, as well as enrichment of interferon and ER-stress response programs (**FDR** ≤ **0.1**, **Figure 4G**). Consistent with these pathway-level changes, approximately two-thirds of SREBF1 transcriptional targets were not up-regulated in 3q-gain cells following mevastatin treatment (**Figure 4H**), indicating attenuated SREBP transcriptional output in the 3q-gain context. Treatment with the SREBP activation inhibitor fatostatin recapitulated genotype-specific suppression of SREBP-responsive genes in 3q-gain cells (**Supplementary Figure S4C**), further supporting attenuated SREBP transcriptional output in the 3q-gain context. Previous proteomic profiling under basal conditions demonstrated that proteins involved in cholesterol biosynthesis, lipid metabolic processes, and SREBP-regulated pathways were reduced in 3q-gain cells relative to WT cells, consistent with attenuated pathway output at the protein level (**Supplementary Figure S4D**) (23). Together, these data indicate that 3q-gain is associated with alterations of lipid storage and cholesterol pathways.

### Chromosome 3q-gain impairs SREBP activation and transcriptional responses to mevalonate pathway inhibition

To understand why 3q-gain cells are unable to tolerate inhibition of the mevalonate pathway, we next examined activation of SREBP1, which coordinates transcriptional responses to sterol depletion (schematic overview in **Figure 5A**). Under sterol-high conditions, SREBP forms a complex with SREBP cleavage–activating protein (SCAP), which binds insulin-induced gene (INSIG) proteins, retaining the SCAP–SREBP complex in the ER and preventing activation (31–34). Under sterol-low conditions, INSIG dissociates from SCAP, allowing SCAP to escort SREBP to the Golgi where it undergoes proteolytic cleavage, releasing a mature transcription factor that induces expression of genes involved in cholesterol and fatty acid biosynthesis including *HMGCR*, *HMGCS1*, and *SCD1* (30,31,33–36). In contrast, sterols or inhibitors of SREBP activation block ER-to-Golgi trafficking and prevent accumulation of the mature nuclear SREBP species. Since SREBP activation requires ER-to-Golgi trafficking mediated by SCAP, we asked whether 3q-gain cells are more dependent on SCAP. CRISPRi-mediated knockdown of *SCAP* with three CRISPRi guides (**Supplementary Figure S5A**) resulted in a greater reduction in viability in 3q-gain cells than in WT cells across all 3 guides tested (*p* = 0.043, 0.029, and 0.049, **Figure 5B**).

**Figure 5.**
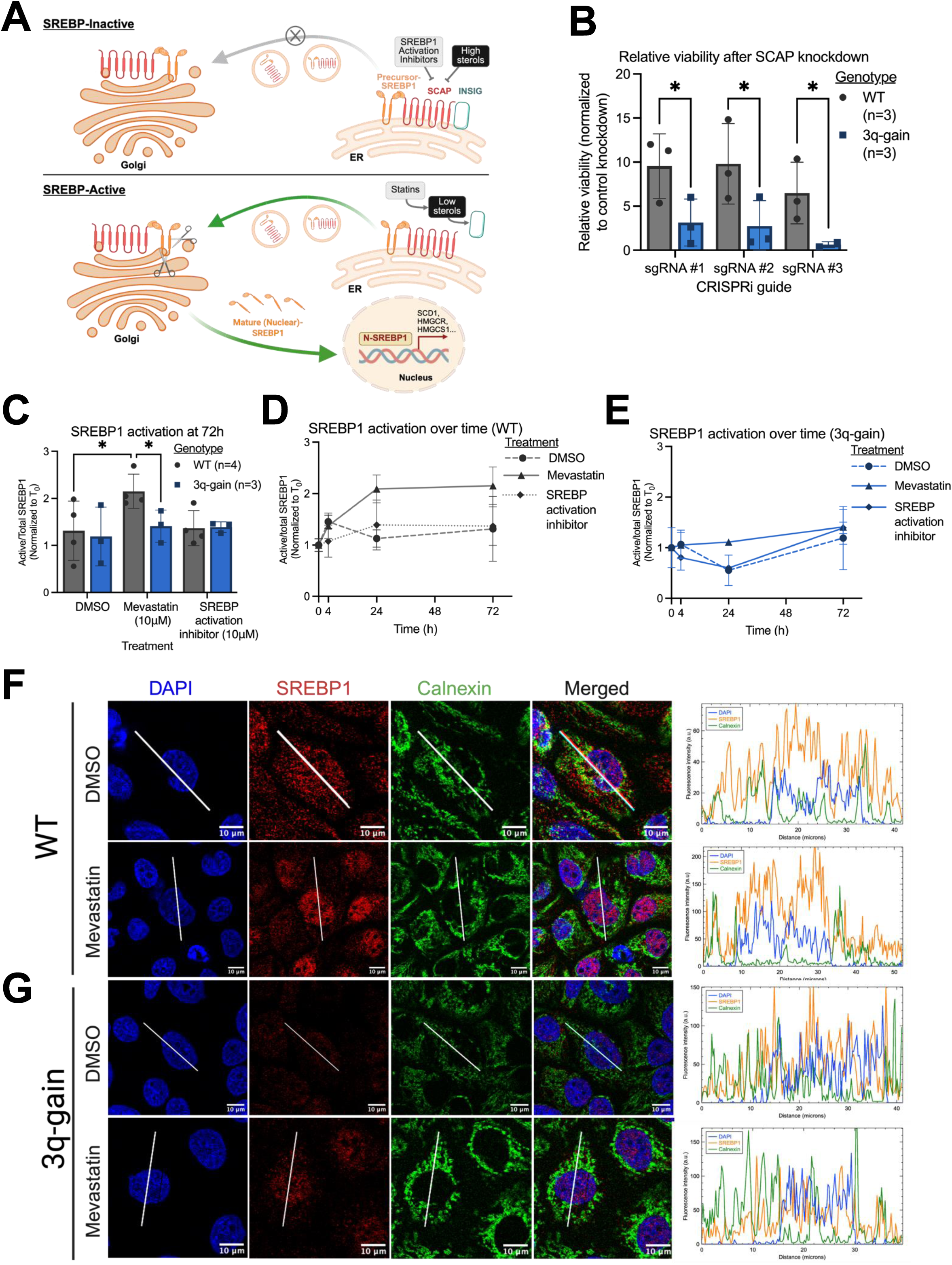
Chromosome 3q-gain impairs activation of SREBP-dependent responses. **(A)** Schematic overview of SREBP1 activation, highlighting ER-to-Golgi trafficking, proteolytic processing, and nuclear translocation in response to sterol depletion, as well as sites of genetic and pharmacologic perturbation used in this study. Created in BioRender. **(B)** Cell number after 5 days of CRISPRi-mediated SCAP knockdown normalized to control sgRNA. **(C)** Quantification of SREBP1 activation at 72 h time point following treatment with DMSO, mevastatin (10 µM), or an SREBP activation inhibitor (10 µM). SREBP1 activation is quantified as the ratio of cleaved (activated) to total SREBP1 and normalized to the corresponding 0 h value. Representative Western blots are shown in **Supplementary Figure S5B-E**. **(D)** Western blot time course of SREBP1 activation in WT clones (*n* = 4) following treatment with DMSO, mevastatin (10 µM), or an SREBP activation inhibitor (10 µM). SREBP1 activation is quantified as in (**C**). Each point represents the average of independent biological replicates. Representative Western blots are shown in **Supplementary Figure S5B-E**. **(E)** Western blot time course of SREBP1 activation in 3q-gain clones (*n* = 3). Each point represents the average of independent biological replicates. Representative Western blots are shown in **Supplementary Figure S5B-E**. **(F-G)** Representative immunofluorescence images showing SREBP1 localization in a WT (**F**) and 3q-gain (**G**) clone treated with DMSO or mevastatin, including DAPI (nuclei), SREBP1, calnexin (ER), merged images, and corresponding line-scan intensity profiles. Images are representative of a single experiment performed across four WT and four 3q-gain clones. Scale bars, 10 µm. Data are shown as mean ± SEM. Dots represent biologically independent clones, quantified as the mean of three technical replicates. Statistical significance was assessed using two-way ANOVA with Tukey’s multiple comparison test for panels **B-C**. *P* value thresholds are defined in Figure 1.

To assess functional activation of the SREBP pathway downstream of sterol sensing, we measured accumulation of the cleaved and mature, transcriptionally active SREBP1 species by western blot. We observed a robust induction of mature SREBP1 in WT cells following 72 h of mevastatin treatment relative to DMSO controls (*p* = 0.018), whereas this response did not occur in 3q-gain cells (**Figure 5C, Supplementary Figure S5B-E**). With mevastatin treatment, SREBP1 activation in WT cells was significantly higher than in 3q-gain cells (*p* = 0.048, **Figure 5C, Supplementary Figure S5B-E**). Treatment with the SREBP activation inhibitor fatostatin suppressed SREBP1 activation in both genotypes (**Figure 5C, Supplementary Figure S5B-E**).

We next assessed accumulation of mature SREBP1 over time following perturbation of sterol availability. In WT cells, mevastatin treatment induced a time-dependent increase in SREBP1 activation over 72 hours, consistent with a sterol-responsive transcriptional program (**Figure 5D, Supplementary Figure S5B-E**). This induction was suppressed by treatment with an SREBP activation inhibitor. In contrast to WT cells, 3q-gain cells exhibited a blunted SREBP1 response to mevastatin, with minimal accumulation of mature SREBP1 across the same time course (**Figure 5E, Supplementary Figure S5B-E**). Consistent with diminished SREBP activation, induction of SREBP target genes was significantly attenuated in 3q-gain cells following mevastatin treatment (FDR ≤ 0.1, **Figure 4G-H**). Expression of lipid biosynthetic enzymes *HMGCS1* and *SCD1* was significantly reduced at both the RNA and protein levels in 3q-gain cells compared to WT cells (**Supplementary Figure S6A-D**), reflecting failure to mount an effective transcriptional response.

We next assessed SREBP1 subcellular localization using immunofluorescence. In vehicle control, SREBP1 exhibited a diffuse distribution spanning the ER and nucleus in both WT and 3q-gain cells, consistent with low-level basal pathway activity (**Figure 5F-G**). Notably, basal SREBP1 signal appeared lower in 3q-gain cells relative to WT under vehicle conditions. In WT cells, mevastatin treatment induced a marked redistribution of SREBP1 toward the nucleus, reflecting increased accumulation of the mature, transcriptionally active form (**Figure 5F**). In contrast, 3q-gain cells displayed overall reduced signal intensity and nuclear enrichment following statin treatment compared to WT cells (**Figure 5G**). Together, these observations indicate that 3q-gain cells display impaired SREBP1 activation of SREBP-dependent transcriptional programs in response to mevalonate pathway inhibition.

### Chromosome 3q-encoded genes modulate sensitivity to mevalonate pathway inhibition

To identify 3q-encoded genes that modulate sensitivity to mevalonate pathway inhibition, we performed a CRISPRi screen targeting genes on chromosome 3q in 3q-gain cells treated with mevastatin or the SREBP activation inhibitor fatostatin (**Supplementary Figure S6E**). In mevastatin-treated cells, the top sensitizing hit was TRK-fused gene (*TFG*), a chromosome 3q-encoded gene implicated in ER-Golgi trafficking and secretory pathway organization (log_2_FC = 2.93, *p* = 8.5 × 10⁻, **Figure 6A**) (37). Independent CRISPRi-mediated knockdown of *TFG* significantly decreased sensitivity to mevastatin in 3q-gain cells compared to control guides (AUC FC: 1.468, *p* = 0.006, **Figure 6B, Supplementary Figure S6F**). In the context of rescue for either drug, we observed a significant enrichment of genes involved in vesicle-mediated transport and membrane trafficking (**Figure 6C, Supplementary Figure S6G-H**). Treatment with brefeldin A, an inhibitor of ER-to-Golgi transport, preferentially reduced viability in 3q-gain cells relative to WT cells (AUC FC: 0.638, *p* = 0.03, **Figure 6D**). Since SREBP activation requires ER-to-Golgi transport for proteolytic processing, these results suggest that altered intracellular trafficking in 3q-gain cells may constrain SREBP activation.

**Figure 6.**
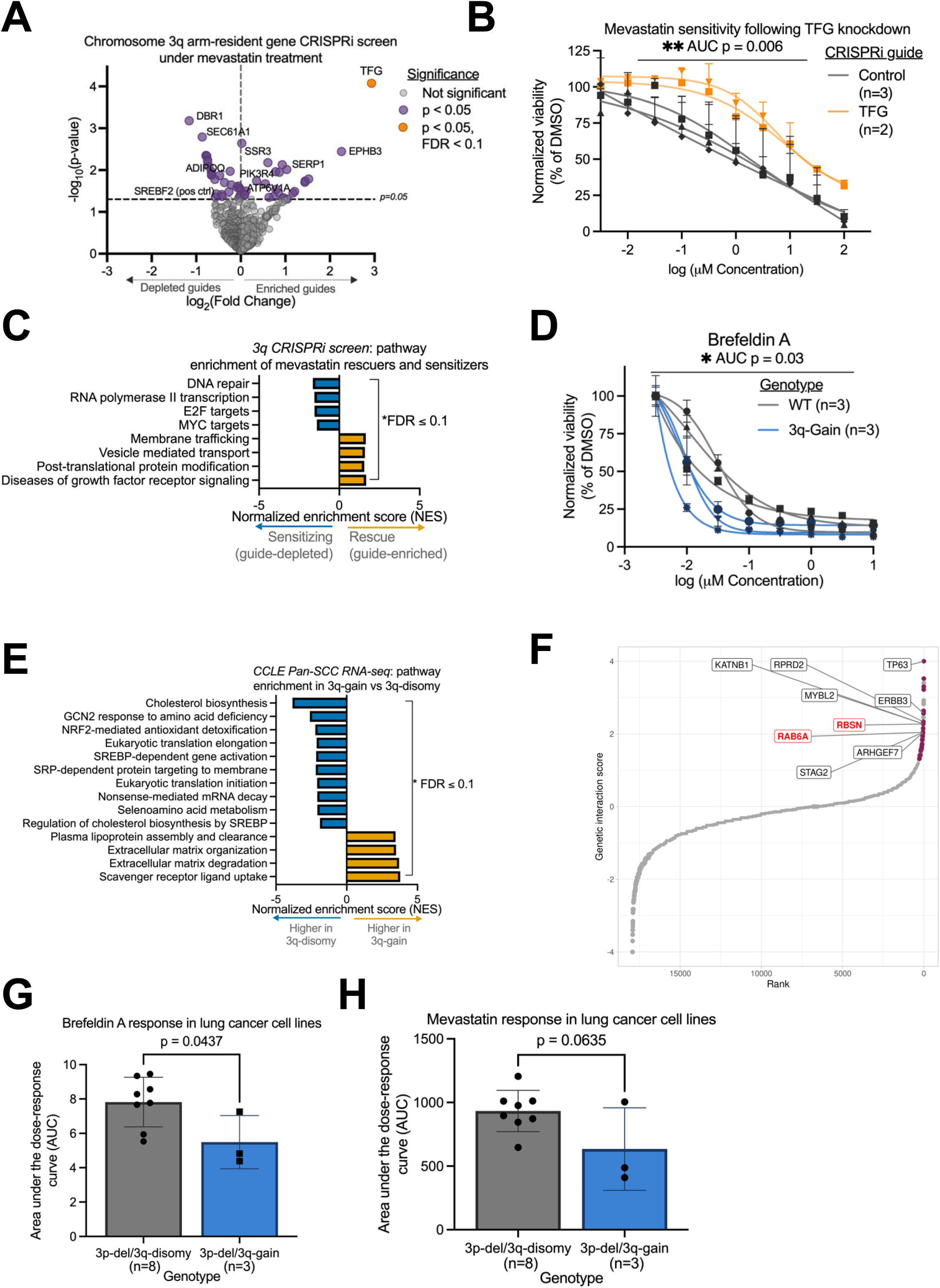
A chromosome 3q-arm CRISPRi screen identifies modifiers of mevalonate pathway sensitivity. **(A)** Volcano plot showing results of the 3q-arm CRISPRi screen under mevastatin treatment. Each point represents an sgRNA, plotted by log₂ fold change and −log₁₀(P value). Highlighted genes represent significantly depleted (sensitizers) or enriched (rescuers) guides (FDR ≤ 0.1). **(B)** Cell viability was measured by CellTiter-Glo 72 h after mevastatin treatment and normalized to DMSO in 3q-gain cells expressing CRISPRi knockdown of *TFG* or a control guide. Each line represents an independent sgRNA (control, *n* = 3; *TFG*, *n* = 2), with each data point reflecting the mean of three technical replicates. Data are averaged across three independent experiments. **(C)** Pre-ranked GSEA of CRISPRi screen hits, showing biological pathways enriched among mevastatin sensitizers (guide-depleted) and rescuers (guide-enriched). NES are shown (FDR ≤ 0.1). **(D)** CellTiter-Glo dose-response analysis of Brefeldin A sensitivity in WT and 3q-gain cells at 72 h, normalized to DMSO. **(E)** Pre-ranked GSEA of RNA-seq data from the Cancer Cell Line Encyclopedia (CCLE) pan-SCC cell lines comparing 3q-gain versus 3q-disomy backgrounds (FDR ≤ 0.1). **(F)** Synthetic lethal interactions of pan-cancer SCC cell lines harboring 3q-gain/3p-deletion predicted *in silico* using GRETTA. Top candidates were ranked by genetic interaction scores. Tier 1 candidates are labeled with red points indicating significance. **(G-H)** Brefeldin A and mevastatin response in lung cancer cell lines stratified by chromosome 3q status in a 3p-deleted background, derived from PRISM drug response data (49). Each dot represents an individual cell line. Data are shown as mean ± SEM. Dots represent biologically independent replicates. Statistical significance was assessed as indicated in the figure; P value thresholds are defined in Figure 1.

### Mevalonate pathway dependence associates with chromosome 3q-gain in squamous cancer models *in vitro, in silico,* and *in vivo*

Some studies have suggested that SREBP promotes proliferation in SCCs (38–46). To assess whether the vulnerabilities identified in isogenic cells are conserved in cancer contexts, we examined transcriptional programs and drug sensitivities associated with 3q-gain in SCC models. In The Cancer Genome Atlas (TCGA) pan–squamous cell carcinoma datasets (9), 3q-gain tumors exhibited depletion of pathways involved in translational initiation and elongation, ER-to-Golgi transport, SRP-dependent protein targeting to membrane, and cellular responses to nutrient deprivation relative to 3q-disomic tumors (FDR ≤ 0.1, **Supplementary Figure S6I**). In SCC cell lines from the Cancer Cell Line Encyclopedia (CCLE) (47), lines with 3q-gain had decreased cholesterol biosynthesis, SREBP-dependent transcription, and SRP-dependent protein targeting to membrane, translation, and stress response pathways compared to 3q-disomy (FDR ≤ 0.1) (**Figure 6E**). Together, these analyses indicate that transcriptional and trafficking-related features associated with 3q-gain are conserved in tumors.

To assess whether 3q-gain is associated with conserved genetic vulnerabilities in cancer, we used GRETTA, a computational framework that leverages the Cancer Dependency Map (DepMap) for *in silico* genetic screening, to predict synthetic lethal (SL) interactions in cancer cell lines harboring 3q-gain (48). In SCC, 3q-gain frequently co-occurs with 3p-deletion. While our epithelial models isolate 3q-gain, several genotype-specific vulnerabilities become more pronounced in cancer models when 3q-gain is present in a 3p-deletion background (13). We therefore evaluated 3q-gain associated dependencies in SCC contexts harboring concurrent 3p-deletion.

We compared 27 SCC carcinoma cell lines with combined 3q-gain and 3p-loss to 6 control cell lines with neutral copy number at both chromosome arms, identifying 46 candidate SL genes whose perturbation preferentially reduced viability in 3q-gain/3p-loss cell lines (adjusted *p* < 0.05; **Figure 6F**; **Supplementary Table S3.1-3.2**). Among these, nine genes were classified as high-confidence Tier 1 candidates. Notably, this set included RBSN and RAB6A, genes with established roles in endosomal and ER-Golgi trafficking, as well as additional candidates involved in intracellular transport and regulatory processes (**Figure 6F**).

We next examined whether these genotype-associated dependencies were evident in pooled drug screening data from lung SCC cell lines harboring 3q-gain on a 3p-deleted background (49). 3p-deletion/3q-gain lung SCC cell lines were significantly more sensitive to disruption of intracellular trafficking, with reduced viability following brefeldin A treatment (*p* = 0.043, **Figure 6G**). We observed a similar trend toward increased sensitivity to mevastatin compared with 3p-loss/3q-disomy controls (*p* = 0.063, **Figure 6H**).

We next wanted to validate these findings in SCC cell lines with and without 3q-gain. Low-pass whole-genome sequencing identified the esophageal SCC cell lines KYSE30 and KYSE520 as 3p-deletion(3p-del)/3q-gain and 3p-del/3q-disomy, respectively (**Supplementary Figure S7A-B**). Consistent with predictions from *in silico* analyses, treatment with mevastatin resulted in significantly greater loss of viability in KYSE30 cells compared to KYSE520 cells (**Figure 7A**), suggesting that 3q-gain confers heightened sensitivity to mevalonate pathway inhibition in a 3p-del SCC. *In vitro* knockdown of HMGCR via CRISPRi also resulted in a significant reduction in viability in 3p-del/3q-gain KYSE30 cells, whereas 3p-del/3q-disomy KYSE520 cells were unaffected (p < 0.0001, **Figure 7B, Supplementary Figure S7C**). HMGCR knockdown in KYSE30 cells also significantly increased apoptotic signaling, measured by Caspase-Glo 3/7 activity (**Supplementary Figure S7D**).

**Figure 7.**
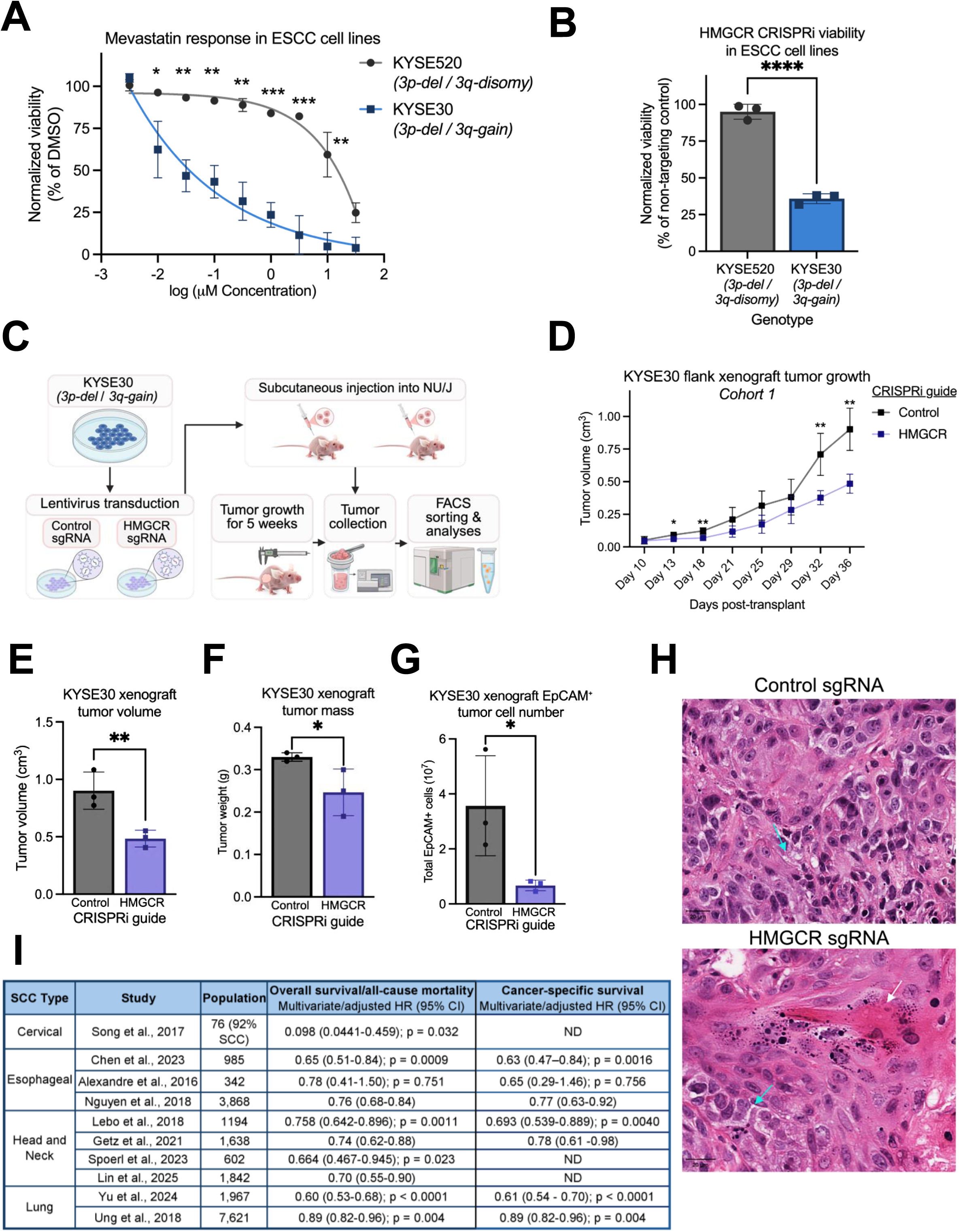
Dependence on mevalonate pathway function associated with 3q-gain extends to squamous cancer models. **(A)** Dose-response curves for mevastatin treatment in esophageal SCC cell lines. Cell viability was measured by CellTiter-Glo 72 h after treatment and normalized to DMSO controls. Data represent mean ± s.d of technical replicates. **(B)** Cell viability following CRISPRi-mediated knockdown of HMGCR in esophageal SCC cell lines. Viability was measured by CellTiter-Glo 5 days after blasticidin selection and normalized to non-targeting control sgRNAs. Each dot represents an independent technical replicate. **(C)** Schematic of *in vivo* xenograft experimental design. KYSE30 cells expressing control or HMGCR-targeting sgRNAs were injected subcutaneously into NU/J mice, and tumor growth was monitored for 5 weeks prior to endpoint collection. Tumors were subsequently dissociated for flow cytometric enrichment of EpCAM-positive epithelial cells and downstream molecular analyses. Created with BioRender. **(D)** Tumor growth curves of KYSE30 xenografts expressing control or HMGCR-targeting sgRNAs. Tumor volume was measured by the caliper and calculated as (length x width^2^)/2. Data represent mean ± s.e.m. of n = 3 mice per group; each point reflects the mean tumor volume of the three mice in the indicated group at each time point. Cohort 1 is shown; data from independent cohorts 2 and 3 are presented in **Supplementary Figure S7E-F**. **(E)** Endpoint tumor volume of KYSE30 xenografts from (**D**). Each dot represents an individual tumor (control sgRNA, n = 3; HMGCR sgRNA, n = 3). **(F)** Endpoint tumor mass of KYSE30 xenografts from (**D**). Each dot represents an individual tumor (control sgRNA, n = 3; HMGCR sgRNA, n = 3). **(G)** Flow cytometric quantification of EPCAM-positive cells from dissociated KYSE30 xenografts from (**D**) at endpoint. Each dot represents an individual tumor (control sgRNA, n = 3; HMGCR sgRNA, n = 3). **(H)** Representative hematoxylin and eosin (H&E) staining of KYSE30 xenograft tumors from Cohort 1 expressing control (**top**) or HMGCR-targeting (**bottom**) sgRNAs. Arrows indicate features of squamous differentiation, including keratinization (white) and intracellular bridges (cyan). Scale bars, 20 µm. **(I)** Summary table of published clinical studies examining associations between statin use and overall survival or all-cause mortality across SCCs. Reported hazard ratios (HRs) reflect multivariate- or covariate-adjusted estimates with corresponding 95% confidence intervals and p-values as reported in the original studies. Statistical significance was assessed by a two-sided Student’s *t*-test. *P* value thresholds are defined in Figure 1.

To assess whether this dependency extends to *in vivo* tumor growth, we generated KYSE30 and KYSE520 cell populations expressing CRISPRi constructs targeting either HMGCR or a non-targeting control and implanted these cells subcutaneously into athymic nude mice (**Figure 7C**). While KYSE520 cells failed to form xenograft tumors under either condition, KYSE30 cells robustly engrafted and formed tumors *in vivo*. Knockdown of HMGCR in KYSE30 cells resulted in a significant reduction in tumor growth over time across three independent *in vivo* cohorts (**Figure 7D**, **Supplementary Figure S7E-F**). Compared to control tumors, HMGCR knockdown significantly reduced tumor volume at endpoint (46% reduction, *p* = 0.007, **Figure 7E**) and tumor mass (25% reduction, *p* = 0.03, **Figure 7F**). At endpoint, tumors were excised and imaged prior to dissociation (**Supplementary Figure S7G**). Tumors were then enzymatically dissociated, and EPCAM-positive cells were isolated by flow cytometry to assess any impact on epithelial tumor cells (**Supplementary Figure S7H**). HMGCR knockdown tumors contained significantly fewer EPCAM-positive epithelial cells (81% reduction relative to controls, *p* = 0.025, **Figure 7G**). RNA isolated from sorted EPCAM-positive cells confirmed sustained HMGCR knockdown *in vivo* by qPCR (**Supplementary Figure S7I**). Histopathologic examination of KYSE30 xenografts revealed features consistent with squamous differentiation, including keratinization and prominent intracellular bridges (**Figure 7H**), confirming the squamous identity of tumors used for *in vivo* assessment. Together, these data demonstrate that 3q-gain is associated with a vulnerability to inhibition of the mevalonate pathway in squamous cancer cells.

Finally, as statin is a widely prescribed drug class, we compiled published clinical studies evaluating associations between statin use and overall survival in SCC (50–59). Across cervical, esophageal, head and neck, and lung SCC cohorts, regardless of HPV status, statin exposure has been recurrently significantly associated with improved overall and cancer-specific survival (**Figure 7I**). Collectively, our findings link 3q-gain with dependency on the mevalonate pathway in epithelial models and squamous cancers.

## Discussion

Unlike many other tumor types, targeted therapeutic strategies have yielded limited improvements in the treatment of SCCs. SCCs, particularly those arising in the upper aerodigestive tract, are not typically driven by oncogenic point mutations (5). Instead, SCCs are characterized by widespread aneuploidy events, with 3q-gain representing one of the most consistent and defining genomic events (4,13,14). The 3q arm encodes a range of putative driver genes, including squamous lineage transcription factors SOX2 and TP63, telomerase RNA component TERC, and proliferative signaling factors such as PIK3CA, PRKCI, ECT2, and MECOM (9,60). CNAs on chromosome 3q are so prevalent that no SCC lines in the Cancer Dependency Map retain complete 3q-disomy, which may explain why SREBF1 does not emerge as a differential vulnerability *in silico*. Our experimental system is uniquely positioned to identify this vulnerability, as we screened immortalized cell lines with and without 3q-gain derived from the putative cell-of-origin for lung SCC, basal epithelial cells (24).

Several genomic and cellular features have previously been associated with differential dependence on the mevalonate pathway. Increased dependence has been reported in p53 mutant or null cells, which is thought to enhance mevalonate pathway activity (61–63). In our model system, immortalization with SV40 functionally suppresses the p53 pathway in both parental (3q-disomy) and derivative (3q-gain) cells providing a controlled context in which p53 signaling is similarly inhibited. Cells with 3q-gain exhibit lower basal SREBP levels compared with 3q-disomy cells, even in the absence of statin treatment. This difference may become apparent only in the context of p53 inhibition, where SREBP levels are otherwise elevated. Directly testing this hypothesis may be challenging, as most SCCs harbor mutations in p53 (9).

Additional cellular states have also been linked to increased dependence on the mevalonate pathway, including mesenchymal phenotypes and the t(4,14) translocation (64). A common feature among sensitive cells is an impaired ability to upregulate SREBP activity following pathway inhibition (65). Our study suggests that 3q-gain contributes to this phenotype: cells harboring 3q-gain fail to mount the typical SREBP response following statin exposure. Consistent with this, 3q-gain cells exhibit reduced lipid droplet abundance, a feature previously associated with increased statin sensitivity (66), reflecting a diminished capacity to buffer perturbations of the mevalonate pathway. The precise mechanism by which 3q-gain attenuates SREBP activation remains unclear; however, our data suggest that altered ER–Golgi trafficking may contribute to impaired SREBP processing and activation.

The potential use of statins in cancer therapy has been investigated for decades (64,67–71). Retrospective pan-cancer analyses have suggested that statin use is associated with an approximately 15% reduction in cancer-specific mortality (72). Squamous-specific clinical cohorts reported in the literature (summarized in **Figure 7I**) tend to demonstrate lower hazard ratios (0.61-0.89) than those observed in broader pan-cancer analyses. Our findings provide a possible mechanistic basis for these observations by linking statin sensitivity to tumors characterized by 3q-gain, a genomic alteration that is particularly prevalent in SCC. A phase-II study evaluating the EGFR inhibitor gefitinib in combination with statins in non–small cell lung cancer reported improved progression-free survival specifically among patients with non-adenocarcinoma histology, of whom 8 of 13 had SCC (73). Although prior studies have examined statin responses in SCC models (74–84), none identified 3q gain as a potential genomic determinant of statin sensitivity.

In summary, our study identifies chromosome 3q-gain as a basis of vulnerability to mevalonate pathway inhibition in SCC. Using a matched cellular model derived from basal epithelial cells, we demonstrate that 3q-gain is associated with impaired SREBP activation in response to statin treatment, potentially due to altered ER–Golgi trafficking. These findings provide a mechanistic framework linking a common aneuploidy event in SCC to metabolic pathway dependence and suggest that statins may have therapeutic value in tumors harboring 3q-gain. Future studies can define the precise molecular drivers within the chromosome 3q region responsible for this phenotype and evaluate whether chromosome 3q status could serve as a predictive biomarker for statin-based therapeutic strategies in SCC.

## Acknowledgments

We thank the members of the A.M.T. laboratory for helpful discussions and feedback on this work. For flow cytometry analyses, we acknowledge support from the Herbert Irving Comprehensive Cancer Center (HICCC) Flow Cytometry Core, supported by National Institutes of Health grant P30CA013696. RNA sequencing of cell lines was performed by the Sulzberger Genome Center at Columbia University. Chemical screening was performed at the Broad Chemical Biology Platform. Genetic screening was performed with support from the Broad Genetics Perturbation Platform (GPP). This work was supported by research funding from Ono Pharmaceutical Co., Ltd., the National Institute of General Medical Sciences (R35GM147287 to A.M.T.), and the National Cancer Institute (pilot funding from P30CA013696 to A.M.T., R01CA273723 to A.M.T., R35CA197568 to M.M.).

## Methods

### Cell Lines and Culture Conditions

Engineered aneuploid cell lines harboring chromosome 3q-gain were previously generated and characterized in XX (female) human lung epithelial cells immortalized with SV40 and TERT (13,24). Briefly, chromosome 3p deletion was induced and a subset of 3p-deleted cells underwent spontaneous adaptation resulting in duplication of the remaining chromosome 3 and acquisition of chromosome 3q-gain. Single-cell cloning was performed to isolate stable 3q-gain populations. For the current study, only validated 3q-gain and chromosome 3 wild-type (WT) clones were used (23). Independent clones were used as biological replicates as indicated in the figure legends. Copy number status was confirmed by low-pass whole-genome sequencing prior to downstream analyses. Cells were maintained at 37°C and 5% CO₂ in Small Airway Growth Medium (Lonza, CC-3118).

KYSE30 and KYSE520 human esophageal squamous cell carcinoma cell lines were obtained from ATCC and cultured at 37°C and 5% CO₂ in RPMI-1640 medium supplemented with 10% fetal bovine serum (FBS) and 1% penicillin-streptomycin. Chromosome 3 copy number status was confirmed by low-pass whole-genome sequencing and ichorCNA (85). HEK293T cells were cultured at 37°C and 5% CO₂ in DMEM supplemented with 10% FBS and 1% penicillin-streptomycin.

### Screens

We screened 5439 arrayed compounds at 2.5 μM in the Broad’s drug repurposing library (27,86) in one line each of 3q-disomy and 3q-gain, with 2 technical replicates per genotype. Z’ was above 0.7 for 3q-disomy and 0.66 for 3q-gain. For initial validation of the top 72 compounds, we generated dose curves in two lines of each genotype. 21 of these compounds passed initial validation were tested with freshly ordered compounds, 3-5 lines of each genotype. Two compounds were considered validated hits, with additional compounds validated beyond the top 72 hit compounds.

For the genome-wide genetic screen, we used the Dolcetto Library Set A (Addgene, 92379) which has 57,050 guides targeting 18,901 genes with 500 unique non-targeting controls. Guides were cloned into an all-in-one vector with dCas9 under an EF-1a promoter and a blasticidin resistance cassette. A pooled lentivirus library and subsequent titering was created as described (25), with 200-250uL of virus per well required for 20-30% efficiency. To achieve 500x coverage, we infected 30 million cells for each of 2 replicates of 2 clones per genotype. Cells were spin-fected (2 hours at 2000 rpm) at 1.25 million cells per well in a 12-well plate with polybrene. One day after infection, cells of the same replicate and clone were pooled and expanded to be at normal density. Two days after this split, 4 μg/mL blasticidin was added and maintained for fourteen days, counting to ensure 30 million cells and then collected for gDNA isolation, PCR (2 per replicate), and sequencing (detailed in Sanson et al., 2021) (25).

For the 3q-specific genetic screen, we created a 3q specific library from Dolcetto Set A, with 1623 guides targeting 541genes encoded on 3q. Along with controls (50 non-targeting guides and 159 guides targeting 53 control genes), we cloned these guides into the same all-in-one vector described above. Lentivirus was produced in HEK293T cells using a calcium phosphate transfection system (Takara). Briefly, 3 × 10 cells were seeded in 10-cm plates and transfected the next day with 4 μg lentiviral sgRNA vector, 4 μg psPAX2 packaging plasmid, and 4 μg pMD2.G (VSV-G) envelope plasmid, with 25 μM chloroquine (Cayman) to enhance transfection efficiency. Viral supernatant was collected 72–96 hours post-transfection, filtered through 0.45 μm filters, validated using Lenti-X GoStix (Takara), and stored at −80 °C. For pooled infection, 3q-gain cells were seeded at 2.5 × 10 cells per 10-cm dish and infected with the pooled library using polybrene-assisted spin-fection. Cells were selected with blasticidin for 5 days. Following selection, an initial T₀ sample was collected for genomic DNA extraction. Cells were then treated with DMSO, 10 μM mevastatin, or 10 μM fatostatin, and samples were collected at multiple time points over approximately two weeks. Genomic DNA was isolated, sgRNA sequences were PCR-amplified and sequenced.

Sequencing reads from genetic screens were processed using MAGeCK. FASTQ files were mapped to the guide RNA library with *mageck count* to generate sgRNA count tables for each sample, including biological replicates (and technical replicates for whole-genome screen). Differential gene analysis between treatment and control samples was performed using *mageck test*, which uses the robust ranking aggregation (RRA) algorithm to assess gene- and sgRNA-level significance. The analysis produced log fold changes (LFC), RRA scores, p-values, and false discovery rates (FDR).

### CRISPR Interference (CRISPRi) Assays

Individual CRISPR interference (CRISPRi) knockdown experiments were performed using sgRNA oligonucleotide sequences listed in **Supplementary Table S4**. Oligos were cloned into the all-in-one backbone containing dCas9 and a blasticidin selection cassette, as described above. Lentivirus was produced in HEK293T cells using the calcium phosphate transfection system (Takara) as described above. Following infection, cells were selected with blasticidin for 5 days to enrich for transduced populations. Cell viability was assessed at multiple time points using Trypan Blue cell counts and CellTiter-Glo assays (Promega) according to the manufacturers’ instructions.

Knockdown efficiency was verified by qPCR. Total RNA was isolated from cells using TRIzol Reagent (Thermo Scientific) according to the manufacturer’s instructions. Reverse transcription was performed using the High-Capacity cDNA Reverse Transcription Kit (Thermo Scientific, 4368814). Quantitative PCR (qPCR) was performed using SYBR Green (Thermo Scientific, 4309155). Relative gene expression was calculated using the 2-ΔΔCt method with normalization to ACTB (β-actin). Primer sequences used for qPCR are provided in **Supplementary Table S5**.

### Cell Viability Assays

Cell lines were seeded in triplicate at 1 x 10^4^ cells per well in 96-well plates in 100μl of medium and allowed to adhere overnight prior to treatment. Cell viability following drug treatment or genetic perturbation was measured using the CellTiter-Glo Luminescent Cell Viability Assay (Promega, G7570; 20 μl). Plates were incubated on a benchtop shaker at room temperature for 10 min before luminescence readings. For CRISPRi assays, cell viability was assessed at multiple time points using CellTiter-Glo as well as Trypan Blue cell counts.

For absolute cell number measurements following drug treatment, cells were seeded and treated as described above, and counts were quantified using automated imaging on an ImageXpress Nano Automated Imaging System (Molecular Devices).

For clonogenic growth assays, cells were seeded at 5 x 10^4^ cells per well in 12-well plates and treated with the indicated drugs for two weeks. Cells were washed with PBS, fixed with 4% formaldehyde, and stained with 0.5% crystal violet in 20% methanol. Plates were air-dried, and bound dye was solubilized using 10% acetic acid. The eluted dye was transferred to 96-well plates in triplicate and absorbance was measured at 590 nm using a Varioskan plate reader (Thermo Scientific).

### Apoptosis Assays

For apoptosis measurements, cells were seeded at 1 x 10^5^ cells per well in 12-well plates and allowed to adhere overnight prior to treatment. Apoptosis was assessed 72 h after treatment using FITC Annexin V (Tonbo, 356409T100) staining for 15-20 min and propidium iodide (MilliporeSigma, P4864) staining followed by flow cytometry on a BD Fortessa. Gating cutoffs for PI- and annexin V-stained samples were determined by comparison to unstained or singly stained samples using FlowJo (BD Biosciences). Early and late apoptotic populations were defined as Annexin V⁺/PI⁻ and Annexin V⁺/PI⁺ cells, respectively. Total apoptosis was defined as the sum of Annexin V⁺/PI⁻ and Annexin V⁺/PI⁺ populations. Apoptosis was also quantified using the Caspase-Glo 3/7 assay (Promega, G8090; 20 μl) 72 h after treatment following plating in triplicate at 1 x 10^4^ cells per well in 96-well plates in 100 μl of medium.

### Drug Treatments and Rescue Experiments

Unless otherwise indicated, drug perturbation assays were analyzed 72 h after treatment. Cells were treated with the indicated concentrations of mevastatin (Cayman Chemical, 10010340), atorvastatin (Cayman Chemical, 10493), pitavastatin (Cayman Chemical, 15414), lovastatin (Cayman Chemical, 10010338), rosuvastatin (Cayman Chemical, 44550), simvastatin (Cayman Chemical, 10010344), fatostatin (Cayman Chemical, 13562), zaragozic acid (Cayman Chemical, 17452), hymeglusin (Cayman Chemical, 11899), RO48-8071 (Cayman Chemical, 10006415), tipifarnib (Cayman Chemical, 11747), brefeldin A (Cayman Chemical, 11861), or 25-hydroxycholesterol (Cayman Chemical, 11097) as specified in each experiment. Compounds were dissolved in DMSO, and vehicle-treated controls were included in all experiments with the final DMSO concentration held constant across conditions.

For dose-response experiments, cells were treated with serial dilutions of the indicated compounds beginning the day after plating. For rescue experiments, mevalonate (MilliporeSigma, M4667) was added concurrently with mevastatin at the indicated concentrations. For drug combination experiments, cells were treated with pairwise drug combinations across dose matrices as indicated. Drug interaction scores were calculated using the Highest Single Agent (HSA) model implemented in SynergyFinder (87).

### Cholesterol and Lipid Quantification Assays

Cells were treated as indicated and intracellular and extracellular cholesterol levels were measured 72 h after treatment using the Cholesterol/Cholesterol Ester-Glo Assay (Promega, J3190) according to the manufacturer’s instructions. Free cholesterol was measured using the detection reagent and total cholesterol was measured following enzymatic conversion of cholesteryl esters to cholesterol using cholesterol esterase. Cholesterol levels were normalized to CellTiter-Glo luminescence to account for differences in cell number as well as baseline measurements at time 0. Cholesterol uptake was measured using the Cholesterol Uptake Assay Kit (Cayman Chemical, 600440) according to the manufacturer’s instructions. Neutral lipid accumulation was assessed using BODIPY 493/503 staining (Thermo Scientific, D3922). Cells were treated as indicated, washed with PBS, and stained with BODIPY as previously described (88). Fluorescence was quantified using flow cytometry on a BD LSRFortessa and analyzed using FlowJo (BD Biosciences).

For lipidomics, lipids were extracted using a methanol/methyl-tert-butyl ether/water (MeOH/MTBE/H_2_O) method containing 0.01% butylated hydroxytoluene (BHT) and SPLASH internal standards (Avanti Research, 330707). Untargeted lipid profiling was performed by liquid chromatography-mass spectrometry (LC-MS) using an ACQUITY I-Class ultra-performance liquid chromatography system (Waters) coupled to a SYNAPT G2-Si HDMS mass spectrometer (Waters) operating in positive and negative electrospray ionization modes. Samples were analyzed in randomized order with duplicates and quality control samples. Data were processed in Progenesis QI (Nonlinear Dynamics) for peak detection and retention-time alignment, normalized to protein concentrations, log-transformed, and analyzed by principal component analysis and partial least squares discriminant analysis. Differential lipids were identified using one-way ANOVA with FDR correction (FDR < 0.05) and annotated according to LIPID MAPS guidelines (89).

### Western Blot

Cells were seeded in 6-well plates at 3 × 10 cells per well and treated as indicated for 72 h, with samples collected at 0 h, 4 h, 24 h, and 72 h. Cells were lysed using RIPA Lysis Buffer System (Santa Cruz), and protein concentration was determined using a Bradford Protein Assay (Cepham Life Sciences). Equal amounts of protein (30 µg) were separated on 4–12% Bis-Tris SDS-PAGE gels (Invitrogen) and transferred to PVDF membranes using the iBlot 2 dry transfer system (Invitrogen). Membranes were blocked and incubated with primary antibodies overnight at 4 °C followed by HRP-conjugated secondary antibodies and detection by chemiluminescence using the SuperSignal West Pico PLUS Chemiluminescent Substrate (Thermo Scientific, 34580). Primary antibodies used were anti-Cofilin (Cell Signaling, 5175), anti-SREBP1 (Proteintech, 14088-1-AP), anti-HMGCS1 (Proteintech, 17643-1-AP), and anti-SCD1 (Proteintech, 28678-1-AP), see **Supplementary Table S6** for ratios for each antibody. HRP-linked anti-rabbit IgG (Cell Signaling, 7074) was used as the secondary antibody. SREBP1 activation was quantified at each timepoint as the ratio of cleaved (mature, ∼68 kDa) SREBP1 to total SREBP1 (cleaved + precursor, ∼125 kDa), based on analysis band intensities using ImageJ, and values were normalized to the 0 h timepoint. Western blots images were generated using SciUGo software.

### Immunofluorescence

Cells were seeded onto glass coverslips in 12-well plates (1 × 10 cells per coverslip) and treated as indicated. Cells were washed with PBS and fixed with 4% paraformaldehyde, followed by permeabilization with 0.1% Triton X-100 in PBS (PBST). Cells were then blocked in 1% BSA and 5% normal donkey serum in PBST prior to antibody staining. Cells were incubated with primary antibodies overnight at 4°C, followed by incubation with fluorophore-conjugated secondary antibodies for 2 h at room temperature. In addition to anti-SREBP1, Calnexin (Thermo Scientific, MA3-027) was used as an endoplasmic reticulum marker. Secondary antibodies include anti-Mouse IgG (AF-488 (Cell Signaling, 4408) and anti-Rabbit IgG (AF-555 (Cell Signaling, 4413). Nuclei were counterstained with DAPI, and coverslips were mounted using Fluoroshield mounting medium (Sigma, F6182). Fluorescence images were acquired using a Leica confocal fluorescence microscope and processed using ImageJ/Fiji.

### Xenograft Studies

All animal experiments were conducted in accordance with protocols approved by the Columbia University Institutional Animal Care and Use Committee and performed in compliance with institutional guidelines for animal welfare. Mice were treatment-naive and maintained under specific pathogen-free conditions in ventilated cages with controlled temperature and humidity on a 12-hour light–dark cycle, with food and water provided ad libitum in the animal care facilities at Columbia University Irving Cancer Research Center.

Female athymic nude mice (NU/J; *Foxn1^nu^*) aged 8 weeks were purchased from The Jackson Laboratory (Stock No. 002019). KYSE30 esophageal squamous cell carcinoma cells expressing sgRNAs targeting HMGCR or a non-targeting control sgRNA were resuspended in a 1:1 mixture of PBS and Matrigel (Corning), and 2.5 × 10 cells were injected subcutaneously into the flanks of mice. Three independent cohorts were injected (n = 3 mice per group per cohort). In cohort 3, one mouse in the control sgRNA group was euthanized prior to study completion due to illness unrelated to tumor burden, and one mouse in the HMGCR sgRNA group was excluded due to development of an internalized tumor; both were omitted from endpoint analyses.

Tumor growth was monitored twice weekly using digital calipers, and tumor volume was calculated as V = (length × width²) / 2. Mice were euthanized after approximately 5 weeks when tumors reached humane endpoints or a maximum volume of approximately 1.5 cm³. Tumors were excised, weighed, imaged, and processed for downstream analyses. Half of each tumor was fixed overnight in 10% neutral-buffered formalin, transferred to 70% ethanol, and processed for paraffin embedding and hematoxylin and eosin (H&E) staining. Histopathologic evaluation was performed by a board-certified pathologist.

The remaining tumor tissue was mechanically dissociated using collagenase P (Sigma, 11249002001) and DNAseI (Sigma, 10104159001) with the gentleMACS Dissociator (Miltenyi Biotec). Cell suspensions were filtered, treated with RBC lysis buffer (Thermo Scientific, 00-4333-57), and stained with APC anti-human CD326 (EpCAM) antibody (BioLegend, 324207), rat anti-mouse CD16/CD32 Fc block (BD Biosciences, 553142), and propidium iodide prior to flow cytometric sorting for human epithelial tumor cells. RNA was isolated from sorted cells using the RNeasy Micro Kit (Qiagen) with 2-mercaptoethanol (Sigma), and qPCR was performed as described above to validate HMGCR knockdown.

### RNA Sequencing and Pathway Enrichment Analysis

Total RNA was extracted using TRIzol (Thermo Scientific, 15596026) followed by purification with the Norgen Total RNA Purification Kit (Norgen Biotek, 17200) according to the manufacturers’ instructions. RNA sequencing libraries were prepared and sequenced on an Illumina platform. Reads were aligned to the human reference genome (GRCh38) and transcript abundances were quantified using kallisto and summarized to gene-level expression using tximport. Gene expression values were normalized as transcripts per million. Differential gene expression analysis was performed using DESeq2 (90). Heatmaps were generated using Morpheus (Broad Institute), and normalized gene expression counts are provided in **Supplementary Table S2**.

Pre-ranked Gene Set Enrichment Analysis (GSEA) was performed using gene lists ranked according to log₂ fold change or screen-derived statistics, as appropriate (91,92). Enriched pathways in Reactome and Gene Ontology gene sets were defined using FDR q < 0.25, as recommended for GSEA.

### DepMap Genetic Dependency Analyses

Squamous cell lines were identified using disease lineage annotations from the DepMap portal (23Q4) (47). Chromosome 3 arm copy-number status was assigned by cross-referencing cell lines with chromosomal arm annotations previously reported (93), defining 3q-gain and 3p-deletion as 3q = 1 and 3p = −1, respectively, and control lines as neutral for both arms (**Supplementary Table S3.1**). Analyses were restricted to microsatellite-stable cell lines. The squamous cell lines were then used as input for the *in silico* genetic dependency analysis, performed using GRETTA (v4.0) (48,94–96) with 3q-gained and 3p-deleted lines as the mutant group and neutral lines as the control. GRETTA used pairwise Mann–Whitney U tests to compare lethality probabilities for all 18,333 genes targeted in the DepMap CRISPR-Cas9 knockout screen between the mutant and control cell line groups. P-values were adjusted for multiple testing using a permutation-based approach (10,000 randomized resamplings). Synthetic lethal candidates were classified into predefined Tier 1–3 categories (94), with Tier 1-2 considered high-confidence (**Supplementary Table S3.2**). Genetic interaction scores calculated by GRETTA were used for ranking and visualization.

Drug sensitivity profiles for Brefeldin A and Mevastatin response in lung cancer cell lines were obtained from the PRISM Repurposing Drug Screen (49). Cell lines were stratified by chromosome 3 status to compare cell lines with 3q-gain versus 3q-disomy in a 3p-deletion background. Area under the curve (AUC) values were used as a measure of drug sensitivity.

## Data Availability

RNA sequencing data from The Cancer Genome Atlas (TCGA) and the Cancer Cell Line Encyclopedia (CCLE) were analyzed to compare 3q-gain and 3q-disomy pan-SCC samples (Campbell et al., 2018; Ghandi et al., 2019). TCGA expression and copy number data is available at https://gdc.cancer.gov/about-data/publications/pancanatlas. CCLE expression, copy number, and chromosomal arm status data, including those used in GRETTA, are available at https://depmap.org/portal/data_page/?tab=overview. RNA-sequencing counts data from our cell lines is in **Supplementary Table S2**, and FASTQ files will be uploaded to SRA upon publication.

### Statistical Analyses

Statistical analyses for the experiments were conducted using GraphPad Prism 10. The graph displays all individual values. Detailed information regarding the tests and corresponding *P*-values is provided in the figures and their respective legends. *P*-values are indicated as follows: * < 0.05, ** < 0.01, *** < 0.001, **** < 0.0001. Unless otherwise indicated, biological replicates correspond to independent cell clones, as specified in figure legends.

### BioRender Licenses

- **Figure 1**. Created in BioRender. Zhakula, N. (2026) https://BioRender.com/bex6ovs
- **Figure 2-3**. Created in BioRender. Zhakula, N. (2026) https://BioRender.com/78ast25
- **Figure 5**. Created in BioRender. Zhakula, N. (2026) https://BioRender.com/tzz4w76
- **Figure 7**. Created in BioRender. Zhakula, N. (2026) https://BioRender.com/0389n5i
- **Supplementary Figure S2**. Created in BioRender. Zhakula, N. (2026) https://BioRender.com/lxu9dfa
- **Supplementary Figure S6**. Created in BioRender. Zhakula, N. (2026) https://BioRender.com/61sy6s5

## Table Legends

**Supplementary Table S1. Quantified lipid species across WT and 3q-gain clones treated with DMSO or mevastatin for 72 h.**

Values represent normalized ion intensities for annotated lipid species measured in WT (*n = 5*) and 3q-gain (*n = 6*) clones following 72 h of treatment with DMSO or mevastatin (10 µM). Columns correspond to individual biological replicates representing independent clones. Lipid species are annotated according to standard lipid nomenclature, where numbers indicate fatty acyl chain length and degree of unsaturation. For lipids containing multiple fatty acyl chains, individual chains are separated by an underscore; for ceramides, the sphingoid base precedes the semicolon and the N-acyl fatty acid chain follows the underscore.

**Supplementary Table S2. Gene-level normalized RNA-seq counts across all WT and 3q-gain clones treated with DMSO, mevastatin, or fatostatin for 72 h.**

Values represent gene-level transcripts per million (TPM) across WT 3q-gain clones treated with DMSO, mevastatin (10 µM), or fatostatin (10 µM). Columns correspond to individual biological replicates representing independent WT (*n = 4*) or 3q-gain clones (*n = 6*).

**Supplementary Table S3.1. DepMap squamous cell carcinoma lines used for GRETTA synthetic lethality analysis.**

List of DepMap squamous cell carcinoma cell lines included in the GRETTA analysis. Cell lines are annotated by disease lineage, subtype, and chromosome 3 arm copy-number status. Chromosome 3q-gain and chromosome 3p-deletion were defined as described in Methods.

**Supplementary Table S3.2. Predicted synthetic lethal candidate genes identified by GRETTA analysis.**

Output of the GRETTA (v4.0) *in silico* genetic interaction analysis identifying candidate synthetic lethal genes associated with chromosome 3q-gain and chromosome 3p-deletion. Columns report median lethality scores in control and mutant groups, log2 fold change calculated from group medians, genetic interaction scores, permutation-adjusted P values, and synthetic lethal classification tiers.

**Supplementary Table S4. CRISPRi sgRNA sequences used in this study.**

Sequences of single guide RNAs (sgRNAs) used for CRISPR interference (CRISPRi) targeting the indicated genes. Non-targeting control sgRNAs were used as negative controls. Oligonucleotide sequences are listed 5′→3′. Applications indicate the experimental systems in which each sgRNA was used.

**Supplementary Table S5. qPCR primer sequences used in this study.**

Forward and reverse primer sequences used for quantitative PCR (qPCR) analysis. Primer sequences are listed in the 5′→3′ orientation.

**Supplementary Table S6. Antibodies used in this study.**

Antibodies used for western blot (WB), immunofluorescence (IF), and flow cytometry. Host species, clonality, vendor, catalog number, and working dilutions for each application are indicated.

## Supplementary Figures

**Supplementary Figure S1.**
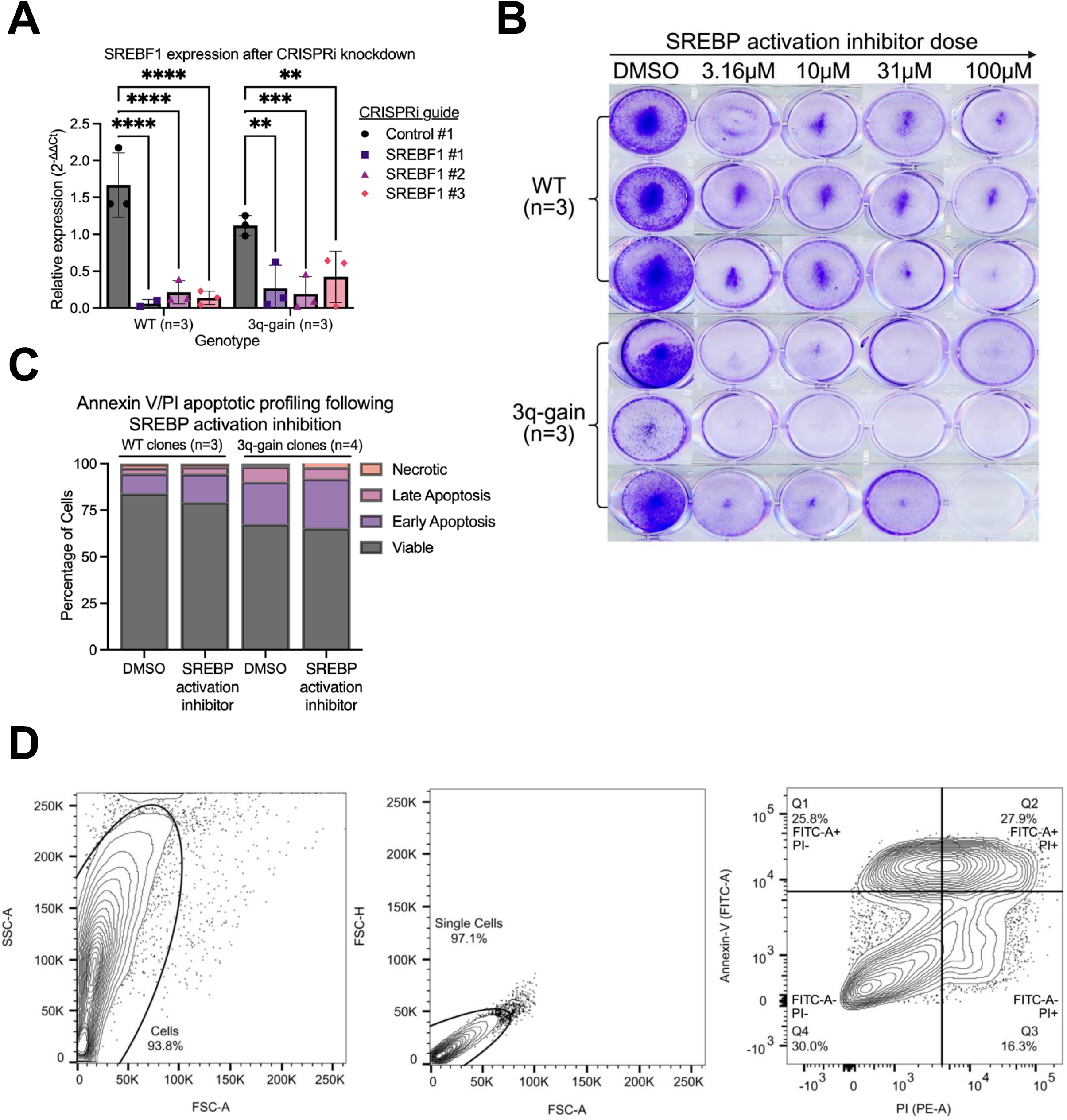
Validation of SREBP1 repression and downstream apoptotic effects following SREBP activation inhibition. **(A)** Relative *SREBF1* mRNA expression following CRISPRi-mediated knockdown using three independent *SREBF1*-targeting sgRNAs (#1–#3) in WT and 3q-gain cells, as measured by qPCR and compared to non-targeting control sgRNA within each genotype. Statistical significance was assessed by two-sided Student’s *t*-test comparing each *SREBF1*-targeting sgRNA to control sgRNA within the same genotype. P values are represented as: *, P < 0.05; **, P < 0.01; ***, P < 0.001; ****, P < 0.0001. **(B)** Clonogenic growth assay of WT and 3q-gain cells treated with increasing concentrations of the SREBP activation inhibitor fatostatin (3.16–100 μM) or DMSO control over two weeks. Rows correspond to independent clones, and columns correspond to independent doses. **(C)** Annexin V/propidium iodide (PI) apoptotic profiling of WT and 3q-gain cell clones following treatment with DMSO or SREBP activation inhibitor for 72 h. Stacked bar plots indicate the percentage of viable (Annexin V^-^/PI^-^), early apoptotic (Annexin V^+^/PI^-^), late apoptotic (Annexin V^+^/PI^+^), and necrotic (Annexin V^-^/PI^+^) populations. Data is consistent with quantified apoptosis measurements shown in Figure 1. **(D)** Representative flow cytometry gating strategy for Annexin V/PI analysis, including forward scatter (FSC) versus side scatter (SSC) gating to exclude debris, singlet discrimination, and quadrant gating to define apoptotic populations. Corresponding quantitative analyses shown in panel **C** and in Figure 1. Unless otherwise indicated, data represent mean ± s.e.m. from *n* biologically independent samples as indicated.

**Supplementary Figure S2.**
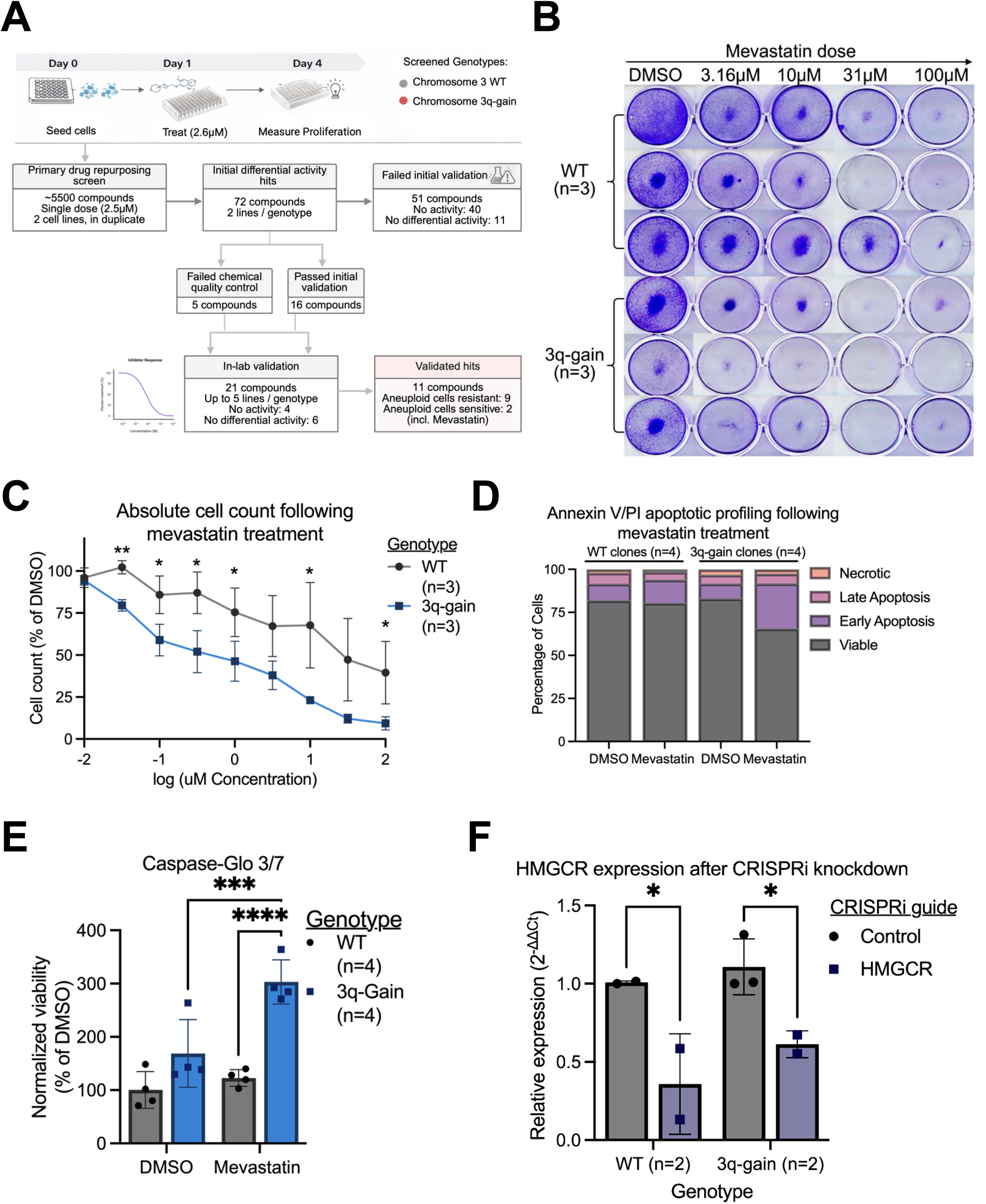
Identification and validation of genotype-specific sensitivity to mevastatin. **(A)** Schematic of the primary drug repurposing screen and validation workflow. WT and 3q-gain cell lines were treated with a single dose of compounds (2.6 μM), and proliferation was assessed after 72 h. Differentially active compounds were subjected to secondary validation and in-lab follow-up, resulting in the identification of aneuploid-specific sensitizers. Created in BioRender. **(B)** Clonogenic growth assay of WT and 3q-gain cells treated with increasing concentrations of mevastatin (3.16–100 μM) or DMSO control over two weeks. Rows correspond to independent clones, and columns correspond to independent doses. **(C)** Absolute cell counts following 72 h of mevastatin treatment across a dose range, normalized to DMSO within each genotype. Statistical significance was assessed by two-sided Student’s *t*-test at individual concentrations. **(D)** Annexin V/PI apoptotic profiling of WT and 3q-gain clones following mevastatin treatment. Stacked bars indicate the percentage of viable, early apoptotic, late apoptotic, and necrotic cells. **(E)** Caspase-Glo 3/7 assay measuring apoptotic activity following mevastatin treatment, normalized to DMSO controls within each genotype. Statistical significance was assessed by two-way ANOVA followed by Tukey’s multiple comparison test. **(F)** qPCR validation of HMGCR knockdown following CRISPRi-mediated targeting in WT and 3q-gain cells. Expression is shown relative to the control guide within each genotype. Statistical significance was assessed by a two-sided Student’s *t*-test. Unless otherwise indicated, data represent mean ± s.e.m. from *n* biologically independent samples as indicated. *P* value thresholds are defined in **Supplementary Figure 1**.

**Supplementary Figure S3.**
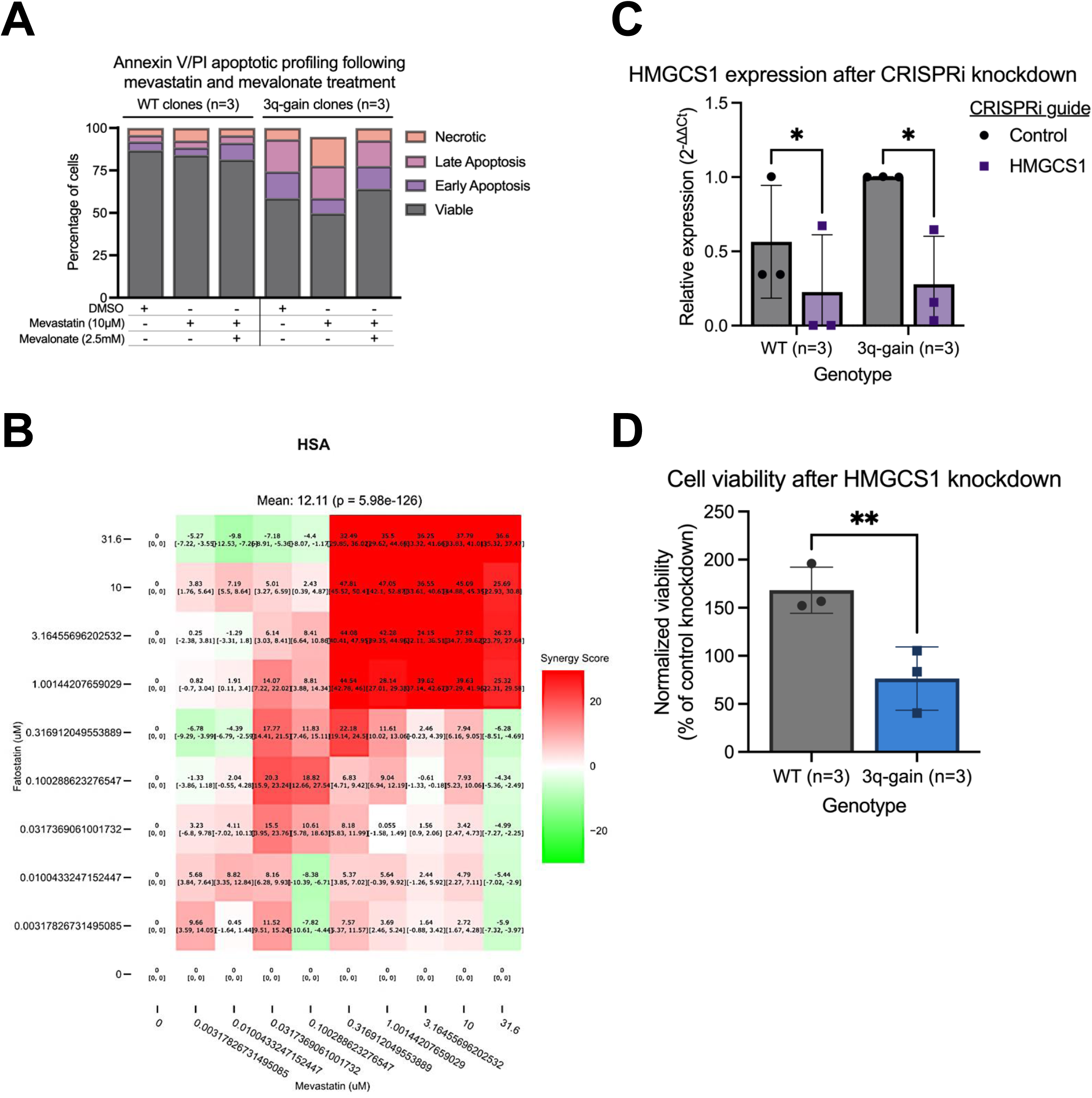
Mevalonate pathway rescue and pharmacologic synergy analyses. **(A)** Annexin V/PI apoptotic profiling of WT and 3q-gain clones following treatment with mevastatin in the presence or absence of mevalonate supplementation. Stacked bars indicate the percentage of viable, early apoptotic, late apoptotic, and necrotic cells. **(B)** Highest Single Agent (HSA) synergy analysis for combined treatment with mevastatin and the SREBP activation inhibitor fatostatin in 3q-gain clones only. The heatmap depicts HSA synergy scores calculated from the mean viability response averaged across three independent 3q-gain clones over the dose matrix, where positive values indicate that the observed combination effect exceeds that of the most effective single agent at matched doses. The mean HSA synergy score across the matrix is shown (mean = 12.11; *p* = 5.98 × 10⁻¹² ). **(C)** qPCR analysis of HMGCS1 expression following CRISPRi-mediated knockdown in WT and 3q-gain cells. Expression is shown relative to the control guide within each genotype. **(D)** Cell viability following CRISPRi-mediated knockdown of HMGCS1 in WT and 3q-gain cells shown in (**C)**, normalized to control knockdown within each genotype. Unless otherwise indicated, data represent mean ± s.e.m. from *n* biologically independent samples as indicated. Statistical significance was assessed by two-sided Student’s *t*-test in panels **C**–**D**. *P* value thresholds are defined in **Supplementary Figure S1.**

**Supplementary Figure S4.**
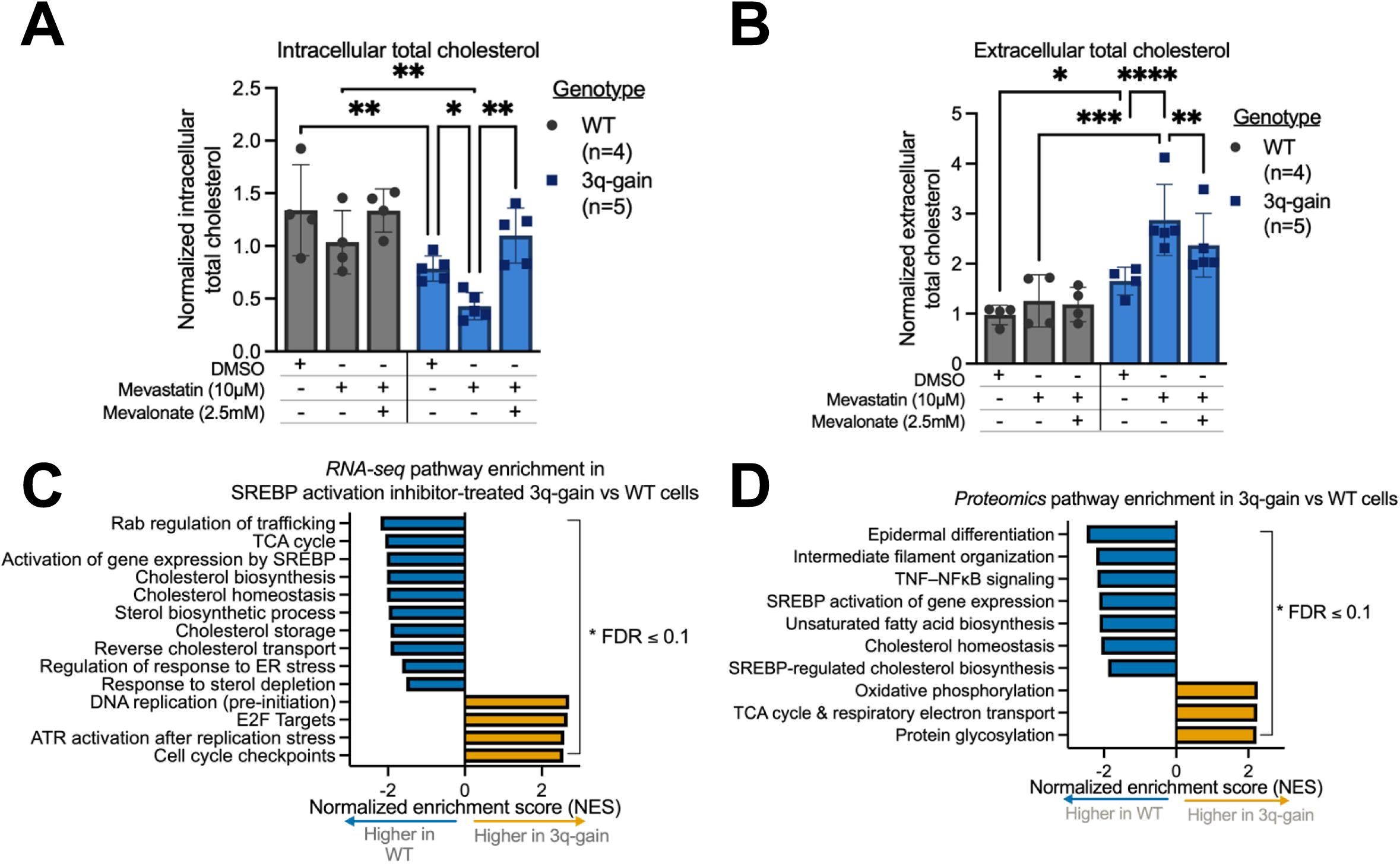
Altered cholesterol homeostasis and SREBP-associated transcriptional programs in 3q-gain cells. **(A)** Intracellular total cholesterol levels in WT and 3q-gain cells following 72 h of treatment with mevastatin in the presence or absence of mevalonate supplementation. Cholesterol levels are quantified with Cholesterol-Glo assay and normalized within each genotype to DMSO-treated controls. **(B)** Extracellular total cholesterol levels measured and normalized as in (**A**). Cholesterol levels are normalized within each genotype to DMSO-treated controls. **(C)** Gene set enrichment analysis (GSEA) of RNA-sequencing (RNA-seq) data comparing 3q-gain versus WT cells treated with the SREBP activation inhibitor fatostatin for 72 h. **(D)** GSEA of proteomics data comparing 3q-gain versus WT cells (23). Unless otherwise indicated, data represent mean ± s.e.m. from *n* biologically independent samples as indicated. Statistical significance was assessed by two-way ANOVA with genotype and treatment as factors, followed by Tukey’s multiple comparison test in panels **A**–**B**. In panels **C**–**D**, pathways with false discovery rate (FDR) ≤ 0.1 are shown, and normalized enrichment scores (NES) indicate relative pathway enrichment in WT (blue) or 3q-gain (yellow) clones. *P* value thresholds are defined in **Supplementary Figure S1.**

**Supplementary Figure S5.**
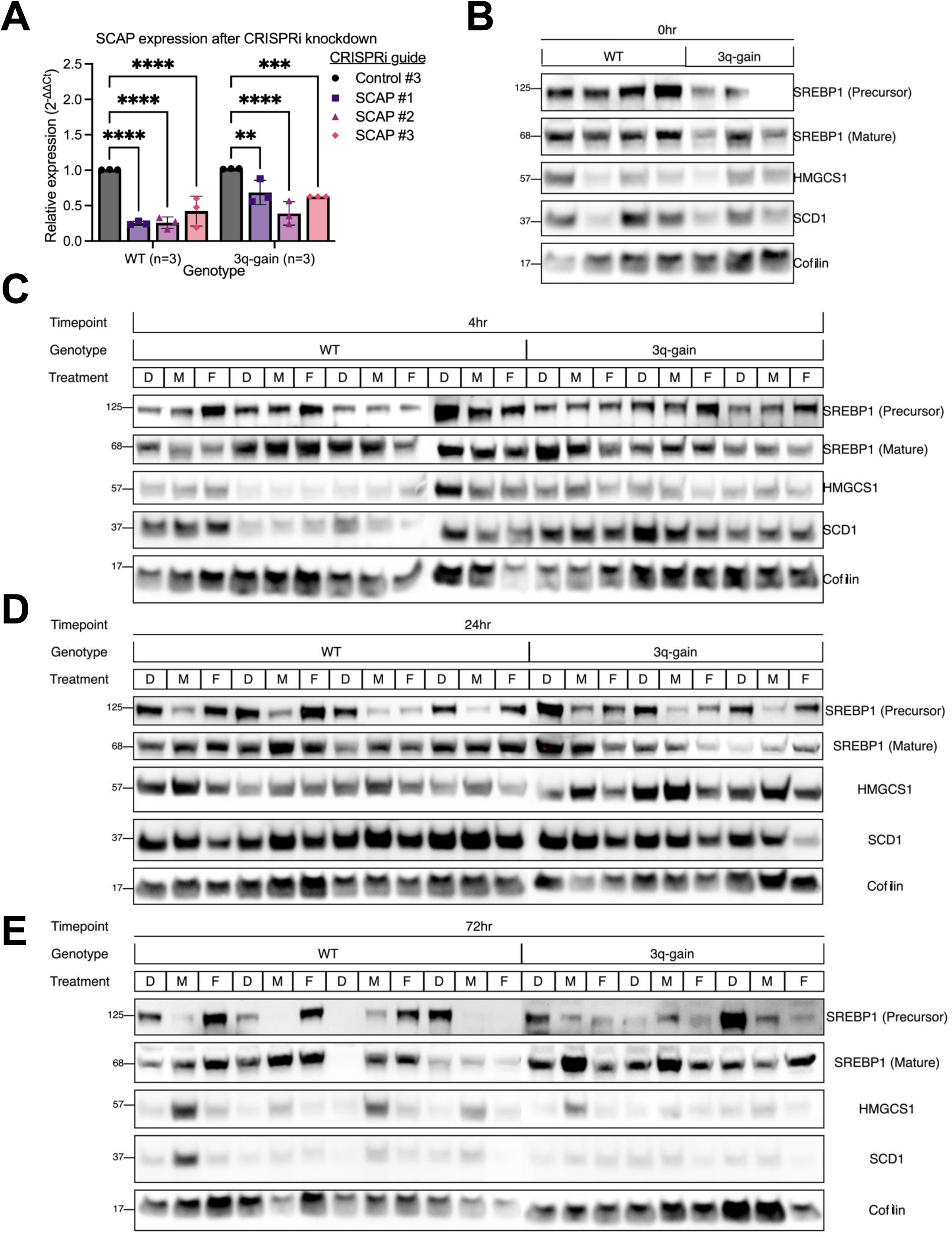
Genetic and pharmacologic perturbation of SREBP activation in WT and 3q-gain cells. **(A)** qPCR analysis of SCAP expression following CRISPRi-mediated knockdown using three independent sgRNAs in WT and 3q-gain cells. Expression is shown relative to the control guide within each genotype. Data represent mean ± s.e.m. (WT, *n* = 3; 3q-gain, *n* = 3). Statistical significance was assessed by one-way ANOVA, followed by Tukey’s multiple comparison test. *P* value thresholds are defined in **Supplementary Figure S1**. **(B-E)** Western blot analysis of SREBP1 precursor and mature forms, and downstream SREBP target proteins (HMGCS1 and SCD1) in WT (*n* = 4) and 3q-gain (*n* = 3) cells across a time course following treatment with DMSO (D), mevastatin (M), or the SREBP activation inhibitor fatostatin (F). Panel **B** shows baseline (0 h) expression, and panels. Panels **C**–**E** show responses following 4 h, 24 h, and 72 h of treatment, respectively. Cofilin is shown as a loading control.

**Supplementary Figure S6.**
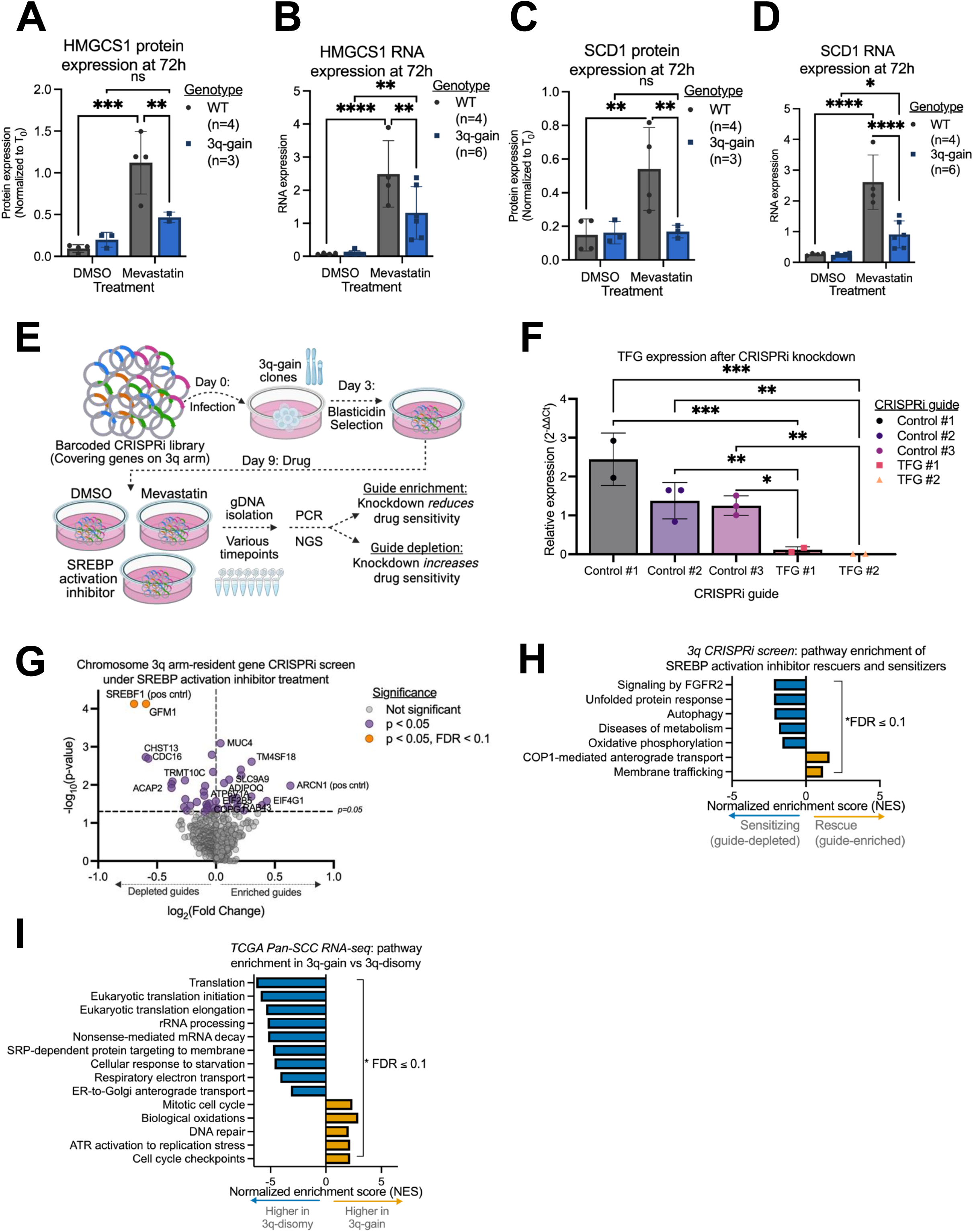
Regulation of mevalonate pathway components and identification of genotype-specific dependencies associated with SREBP perturbation and 3q-gain. (A-D) Expression of HMGCS1 (**A**–**B**) and SCD1 (**C**–**D**) following 72 h of mevastatin treatment in WT and 3q-gain cells. (**A**) HMGSC1 protein expression and (**B**) HMGCS1 RNA expression; (**C**) SCD1 protein expression and (**D**) SCD1 RNA expression. Protein expression in panels **A** and **C** were quantified from the 72 h time course shown in Figure 5C–**E** and in representative western blots in **Supplementary Figure S5B**–**E.** RNA expression in panels **B** and **D** were derived from the RNA-sequencing data at the same timepoint, shown in Figure 4F-H and **Supplementary Figure S4C**. All values were compared to DMSO-treated controls within each genotype. Data represent mean ± s.e.m. from *n* biologically independent samples as indicated. Statistical significance was assessed by two-way ANOVA with genotype and treatment as factors, followed by Tukey’s multiple comparison test. **(E)** Schematic of the chromosome 3q-focused CRISPRi screen performed in 3q-gain clones. A barcoded CRISPRi library targeting genes located on chromosome 3q was introduced into five 3q-gain clones, followed by blasticidin selection and treatment with mevastatin, the SREBP activation inhibitor fatostatin, or DMSO. Guide representation was quantified by next-generation sequencing at multiple time points to identify guides whose depletion or enrichment altered drug sensitivity. Created in BioRender. **(F)** qPCR analysis of TFG expression in 3q-gain clones following CRISPRi-mediated knockdown using two independent sgRNAs, with three non-targeting control guides shown for comparison. Expression is shown relative to control guides. Data represent mean ± s.e.m. Statistical significance was assessed by one-way ANOVA followed by Tukey’s multiple comparison test. **(G)** Volcano plot summarizing results of the chromosome 3q CRISPRi screen under SREBP activation inhibitor treatment. Guides are plotted by log₂ fold change versus −log₁₀(*p*) value. Genes meeting significance thresholds (*p* < 0.05 and FDR < 0.1) are highlighted, including positive controls. **(H)** Pathway enrichment analysis of sensitizing (guide-depleted) and rescuing (guide-enriched) hits identified in the chromosome 3q CRISPRi screen performed under treatment with the SREBP activation inhibitor fatostatin (panel **F**). NES indicates sensitizing (negative NES) or rescuing (positive NES) effects. **(I)** GSEA of TCGA pan-SCC RNA-seq data comparing 3q-gain versus 3q-disomic tumors. Pathways with false discovery rate (FDR) ≤ 0.1 are shown, and NES indicate relative pathway enrichment in 3q-disomic (blue) or 3q-gain (yellow) tumors.

**Supplementary Figure S7.**
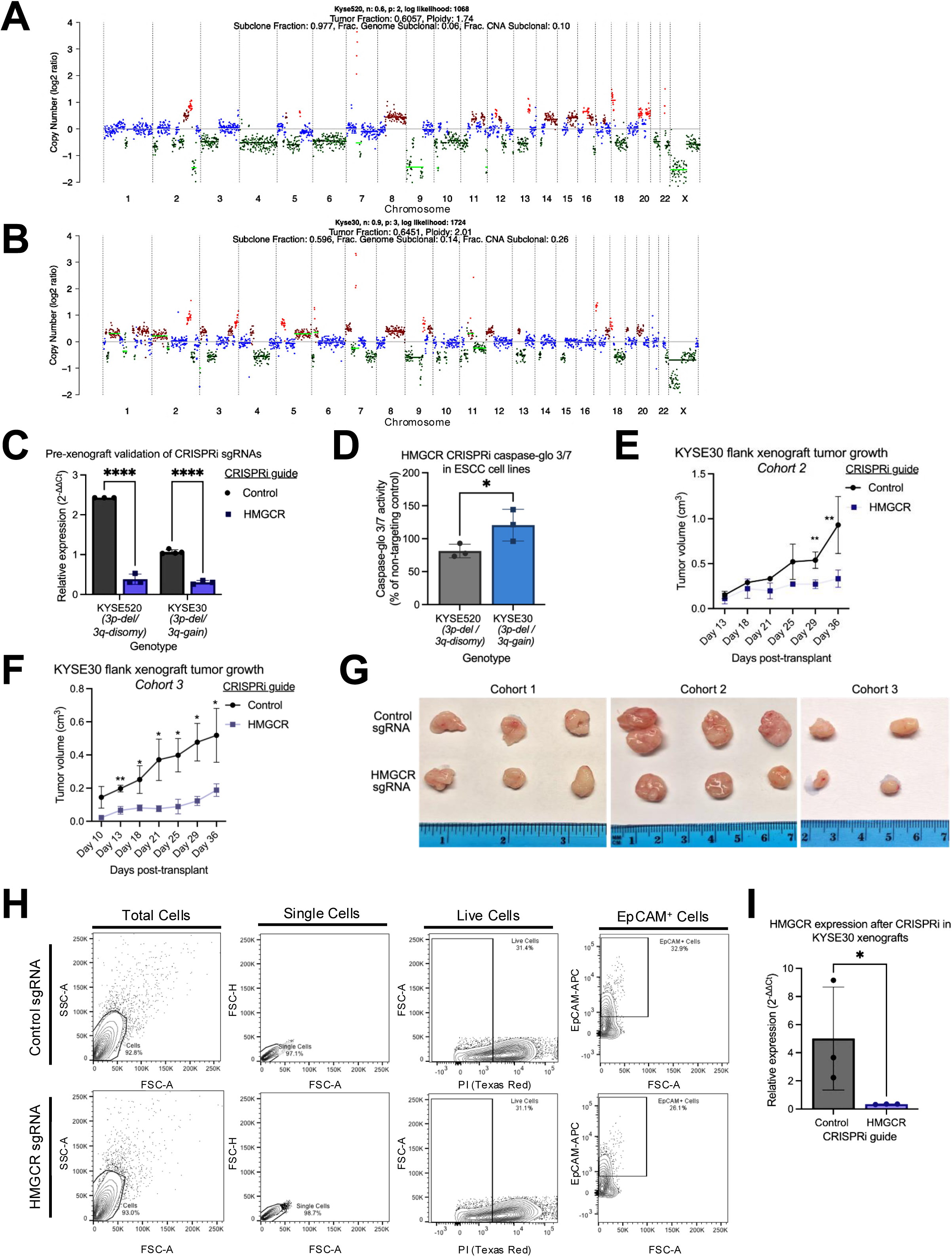
Genotype-specific dependency on HMGCR *in vitro* and in vivo. (A-B) Copy-number profiles of (**A**) KYSE520 (3p-deletion/3q-disomic) and (**B**) KYSE30 (3p-deletion/3q-gain) cells inferred from low-pass whole-genome sequencing, showing relative copy number across the genome. **(C)** qPCR of CRISPRi sgRNAs targeting HMGCR in KYSE520 and KYSE30 cells. Expression is shown relative to non-targeting control guides. Statistical significance was assessed by a two-sided paired Student’s *t-*test. **(D)** Caspase-Glo 3/7 activity following CRISPRi-mediated knockdown of HMGCR in KYSE520 and KYSE30 cells, normalized to non-targeting control guides within each genotype. Statistical significance was assessed by a two-sided Student’s *t*-test. **(E–F)** Tumor growth curves for flank xenografts derived from KYSE30 cells expressing control or HMGCR-targeting CRISPRi guides are shown for cohort 2 (**E**; *n* = 3 control, *n* = 3 HMGCR) and cohort 3 (**F**; *n* = 2 control, *n* = 2 HMGCR). Cohort 1 is shown in Figure 7D-H (*n* = 3 control, *n* = 3 HMGCR). Tumor volume was measured by serial caliper measurements. Statistical significance was assessed by two-sided Student’s *t*-test at individual time points. **(G)** Representative images of excised KYSE30 xenograft tumors from three independent cohorts (cohorts 1-3) expressing control or HMGCR-targeting CRISPRi guides at endpoint. Cohort sizes were *n* = 3 control and *n* = 3 HMGCR-targeting tumors (cohort 1), *n* = 3 control and *n* = 3 HMGCR-targeting tumors (cohort 2), and *n* = 2 control and *n* = 2 HMGCR-targeting tumors (cohort 3). **(H)** Flow cytometry gating strategy used to isolate EPCAM-positive epithelial tumor cells from dissociated KYSE30 xenograft tumors. Cells were first gated based on FSC and SSC to exclude debris, followed by exclusion of dead cells using PI (PE–Texas Red channel). Viable cells were then further gated for EPCAM-positive and EPCAM-negative populations using APC fluorescence. **(I)** qPCR analysis of HMGCR expression in EPCAM-positive cells isolated from KYSE30 xenografts from cohort 1, confirming sustained knockdown *in vivo*. Expression is shown relative to the control guide. Statistical significance was assessed by a two-sided paired Student’s *t*-test. Unless otherwise indicated, data represent mean ± s.e.m.

## Notes

**Conflict of Interest:** A.M.T. and M.M. received research support from Ono Pharmaceuticals for this work. M.M., A.M.T., and N.Z.K. are inventors of a patent application related to this work (WO/2025/255338). A.M.T. is an equity holder of Karyoverse. M.M. is an equity holder of, consultant for, and receives research support from Bayer, and receives patent royalties from Bayer and Labcorp. M.M. is also an equity holder of and consultant for Delve Bio, and an equity holder of Karyoverse and Isabl. M.M. has previously received research support from Janssen and Ono Pharmaceutical and has held equity and consulting roles with Foundation Medicine. B.H. has received institutional research funding from Genentech-Roche, NexImmune, and Johnson & Johnson; speaker honoraria from OncLive; and advisory board fees from AstraZeneca, Ideaya Biosciences, Jazz Pharmaceuticals, Sorrento Therapeutics, Regeneron, Bristol Myers Squibb, Genentech-Roche, Synthekine, Boehringer Ingelheim, and Bayer. All other authors declare no potential conflicts of interest.

### Competing Interest Statement

A.M.T. and M.M. received research support from Ono Pharmaceuticals for this work. M.M., A.M.T., and N.Z.K. are inventors of a patent application related to this work (WO/2025/255338). A.M.T. is an equity holder of Karyoverse. M.M. is an equity holder of, consultant for, and receives research support from Bayer, and receives patent royalties from Bayer and Labcorp. M.M. is also an equity holder of and consultant for Delve Bio, and an equity holder of Karyoverse and Isabl. M.M. has previously received research support from Janssen and Ono Pharmaceutical and has held equity and consulting roles with Foundation Medicine. B.H. has received institutional research funding from Genentech-Roche, NexImmune, and Johnson & Johnson; speaker honoraria from OncLive; and advisory board fees from AstraZeneca, Ideaya Biosciences, Jazz Pharmaceuticals, Sorrento Therapeutics, Regeneron, Bristol Myers Squibb, Genentech-Roche, Synthekine, Boehringer Ingelheim, and Bayer. All other authors declare no potential conflicts of interest.

